# A circuit mechanism for irrationalities in decision-making and NMDA receptor hypofunction: behaviour, computational modelling, and pharmacology

**DOI:** 10.1101/826214

**Authors:** Sean E. Cavanagh, Norman H. Lam, John D. Murray, Laurence T. Hunt, Steven W. Kennerley

## Abstract

Decision-making biases can be systematic features of normal behaviour, or deficits underlying neuropsychiatric symptoms. We used behavioural psychophysics, spiking-circuit modelling and pharmacological manipulations to explore decision-making biases in health and disease. Monkeys performed an evidence integration task in which they showed a pro-variance bias (PVB): a preference to choose options with more variable evidence. The PVB was also present in a spiking circuit model, revealing a neural mechanism for this behaviour. Because NMDA receptor (NMDA-R) hypofunction is a leading hypothesis for neuropathology in schizophrenia, we simulated behavioural effects of NMDA-R hypofunction onto either excitatory or inhibitory neurons in the model. These were tested experimentally using the NMDA-R antagonist ketamine, yielding changes in decision-making consistent with lowered cortical excitation/inhibition balance from NMDA-R hypofunction onto excitatory neurons. These results provide a circuit-level mechanism that bridges across explanatory scales, from the synaptic to the behavioural, in neuropsychiatric disorders where decision-making biases are prominent.

**Significance:** People can make apparently irrational decisions because of underlying features in their decision circuitry. Deficits in the same neural circuits may also underlie debilitating cognitive symptoms of neuropsychiatric patients. Here, we reveal a neural circuit mechanism explaining an irrationality frequently observed in healthy humans making binary choices – the pro-variance bias. Our circuit model could be perturbed by introducing deficits in either excitatory or inhibitory neuron function. These two perturbations made specific, dissociable predictions for the types of irrational decisionmaking behaviour produced. We used the NMDA-R antagonist ketamine, an experimental model for schizophrenia, to test if these predictions were relevant to neuropsychiatric pathophysiology. The results were consistent with impaired excitatory neuron function, providing important new insights into the pathophysiology of schizophrenia.

## Introduction

Schizophrenia is a debilitating neuropsychiatric disorder, associated with prominent deficits in cognitive function^1–3^. Despite being the focus of intensive research, the neural bases of its symptomatology remain poorly understood. Our current understanding of the pathophysiology of schizophrenia mainly focuses on disruptions at the synaptic level. One line of investigations implicates N-methyl-D-aspartate receptor (NMDA-R) dysfunction^4–6^, and NMDA-R antagonists have been used as a pharmacological model of schizophrenia. When administered to healthy volunteers, they transiently reproduce multiple aspects of the symptoms of schizophrenia, especially cognitive deficits^7–9^. One interpretation of these observations is that NMDA-R hypofunction causes an imbalance of excitation and inhibition in cortical circuits^5,10,11^. However, linking these pathophysiological mechanisms to the cognitive impairment observed in patients has proved challenging.

One difficulty is to carefully isolate which cognitive computations underlie neuropsychiatric symptoms. Working memory deficits in patients with schizophrenia have been well-characterised, which has facilitated preclinical research providing insights into potential pathophysiological mechanisms^2,12^. However, whether these working memory deficits reflect a more general impairment in other temporally extended cognitive processes in the symptomatology of schizophrenia remains an open question. One closely related cognitive process is evidence accumulation – the decision process whereby multiple samples of information are combined over time to form a categorical choice^13^. It has been extensively studied using the random-dot motion (RDM) task, where subjects must decide the net direction of a moving dots stimulus^13,14^. Patients with schizophrenia have impaired perceptual discrimination on the RDM task^15–17^, but the precise nature of this decision-making deficit is unclear. Previous studies have attributed it to an impaired representation of the sensory evidence in visual cortex^15,18^, yet circuit-level alterations affecting visual cortex are likely also present in downstream cortical association areas involved in evidence accumulation and decision-making. It is therefore important to characterise precisely whether and how the underlying process of evidence accumulation may be affected in schizophrenia.

Recent research has advanced our understanding of how such evidence accumulation decisions are made in the healthy brain. Of particular relevance to psychiatric research, it has been possible to disentangle systematic biases in decision-making and reveal the mechanisms through which they occur. For instance, when choosing between two series of bars with distinct heights, people have a preference to choose the option where evidence is more broadly distributed across samples^19,20^. Although this “pro-variance bias” may appear irrational, and would not be captured by many normative decision-making models, it becomes the optimal strategy when the accumulation process is contaminated by noise^19^. These behaviours have presently been well-characterised using algorithmic level descriptions of decision formation. By extending this approach to psychiatric research, new insights could be gained into the decision making deficits in schizophrenia. However, in order to understand how these decision biases might be affected by NMDA-R hypofunction, a more mechanistic explanation is needed.

An influential technique used to investigate evidence accumulation at the mechanistic level has been biophysically grounded computational modelling of cortical circuits^21–23^. Through strong recurrent connections between similarly tuned pyramidal neurons, and NMDA-R mediated synaptic transmission, these circuits can facilitate the integration of evidence across long timescales. Crucially, these neural circuit models bridge synaptic and behavioural levels of understanding, by predicting both choices and their underlying neural activity. These predictions reproduce key experimental phenomena, mirroring the behavioural and neurophysiological data recorded from macaque monkeys performing the RDM task^21,24^. Whether neural circuit models can provide a mechanistic implementation of the pro-variance bias, and other irrational aspects of evidence accumulation, is currently unknown. Circuit models also present a promising avenue to address the challenges of neuropsychiatric research due to their biophysically detailed mechanisms. By perturbing the circuit model at the synaptic level, specific behavioural and neural predictions can be made. Relevant to schizophrenia, NMDA-R hypofunction can be introduced to alter the balance between excitation and inhibition (E/I balance)^25^. Recent studies have used NMDA-R antagonists to validate model predictions during working memory tasks^25,26^. While NMDA-R antagonists have been tested during various decision-making tasks^27,28^, the role of the NMDA-R in shaping the temporal process of evidence accumulation has not been characterised experimentally.

Here we used a psychophysical behavioural task in macaque monkeys, in combination with spiking cortical circuit modelling and pharmacological manipulations, to gain new insights into decisionmaking biases in both health and disease. We trained two subjects to perform a challenging decisionmaking task requiring the combination of multiple samples of information with distinct magnitudes. Replicating observations from humans, monkeys showed a pro-variance bias. The pro-variance bias was also present in the spiking circuit model, revealing an explanation of how it may arise through neural dynamics. We then investigated the effects of NMDA-R hypofunction in the circuit model, by perturbing NMDA-R function at distinct synaptic sites. Perturbations could either raise or lower the E/I ratio, with each effect making dissociable predictions for evidence accumulation behaviour. These model predictions were tested experimentally by administering monkeys with a subanaesthetic dose of the NMDA-R antagonist ketamine (0.5mg/kg, intramuscular injection). Ketamine produced decisionmaking deficits consistent with a lowering of the cortical E/I ratio.

## Results

To study evidence accumulation behaviour in non-human primates, we developed a novel two-alternative perceptual decision-making task (**Fig 1a**). Subjects were presented with two series of eight bars (evidence samples), one on either side of central fixation. Their task was to decide which evidence stream had the higher/lower average bar height, and indicate their choice contingent on a contextual cue shown at the start of the trial. The individual evidence samples were drawn from Gaussian distributions, which could have different variances for different options (**Fig 1b**). This task design had several advantages over evidence accumulation paradigms previously employed with animal subjects. Subjects were given eight evidence samples with distinct magnitudes (**Fig 1c**) – encouraging a temporal integration decision-making strategy. Precise experimental control of the stimuli facilitated analytical approaches probing the influence of evidence variability and time course on choice, and allowed us to design specific trials that attempted to induce systematic irrationalities in choice behaviour.

**Figure 1.**
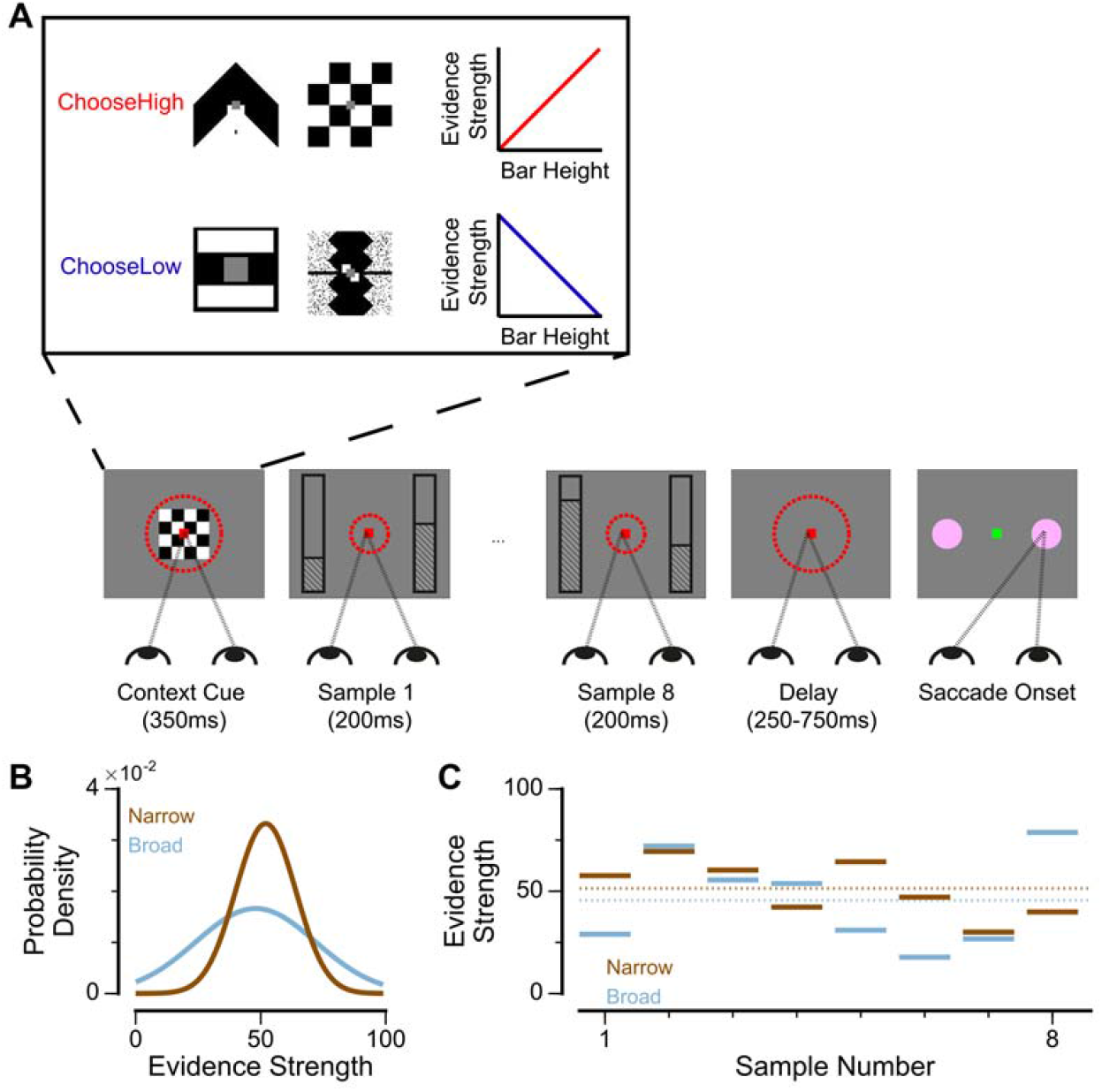
An evidence-varying decision-making task for macaque monkeys. (**A**) Task design. Two streams of stimuli were presented to a monkey, both of which consisted of a sequence of eight samples of bars of varying heights. Depending on the contextual cue shown at the start of the trial, the monkey had to report the stream with either higher or lower mean height. On correct trials, the monkey was rewarded proportionally to the mean evidence for the correct stream; incorrect trials were not rewarded. The monkey was required to fixate centrally while the evidence was presented, indicated by the dashed red fixation zone (not visible to subject). (**B**) Generating process of each stimulus stream. The generating mean for each trial was chosen from a uniform distribution (see Methods), while the generating standard deviation was 12 and 24 for the narrow (brown) and broad (blue) streams respectively. (**C**) Example Trial. The bar heights in both streams varied over time. The dotted lines illustrate the mean of the eight stimuli for the narrow/broad streams. In this example, the narrow stream has a higher mean evidence strength, so is the correct choice. The narrow/broad streams are randomly assigned to the left/right options on different trials; in the example trial shown here (A and C), the narrow stream is assigned to the right option, the broad stream is assigned to the left option.

Two monkeys (*Macaca mulatta*) completed 29,726 trials (Monkey A: 10,748; Monkey H: 18,978). Despite the challenging nature of the task, subjects were able to perform it with high accuracy (**Fig 2a-b**). The precise control of the discrete stimuli allowed us to evaluate the impact of evidence presented at each time point on the final behavioural choice, via logistic regression (see **Methods**). Stimuli presented at a time point with a larger regression coefficient have a strong impact on the choice, relative to time points with smaller coefficients. We found that the subjects utilised all eight stimuli throughout the trial to inform their decision, and demonstrated a primacy bias such that early stimuli have stronger temporal weights than later stimuli (**Fig 2c-d**). A primacy bias has been reported in prior studies in monkeys, and is consistent with a decision-making strategy of bounded evidence integration^29–31^. As it was clear both monkeys could accurately perform the task, all subsequent figures are presented with data collapsed across subjects for conciseness, but results separated by subjects are consistent (**Supplementary Material**).

**Figure 2.**
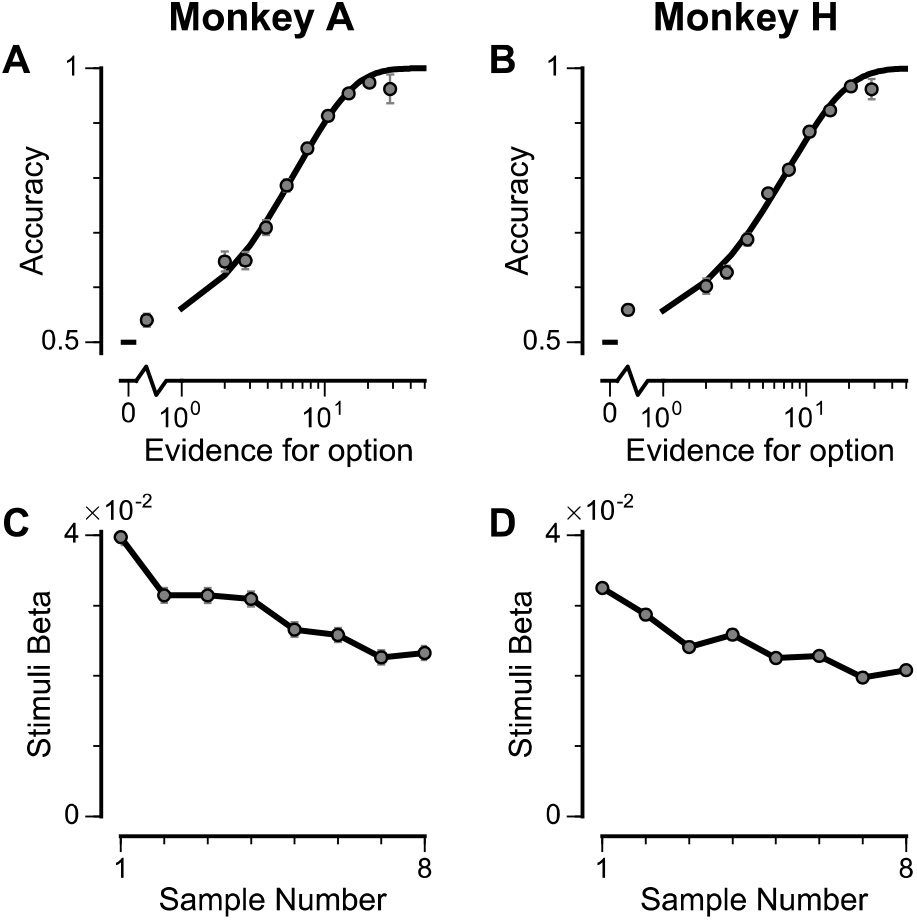
Subjects use evidence presented throughout the trial to guide their choices. (**A-B**) Choice accuracy plotted as a function of the amount of evidence in favour of the best option. Lines are a psychometric fit to the data. (**C-D**) Logistic regression coefficients reveal the contribution (weight) of all eight stimuli on subjects’ choices (see Methods). Although subjects used all eight stimuli to guide their choices, they weighed the initially presented evidence more strongly. All errorbars indicate the standard error.

We next probed the influence of evidence variability on choice. We designed specific choice options with different levels of standard deviation across samples in an attempt to replicate the pro-variance bias previously reported for human subjects (see **Methods**)^19,20^. On each trial, one option was allocated a narrow distribution of bar heights, and the other a broad distribution. In different conditions, either the broad or narrow stimuli stream could be the correct choice (*‘Broad Correct’ Trials or ‘Narrow Correct’ Trials*), or there could be no clear correct answer (*‘Ambiguous’ Trials*) (**Fig 3a, Supplementary Fig. 1**). If subjects chose optimally, and only the mean bar height influenced their choice, their accuracy would be the same in *‘Broad Correct’ and ‘Narrow Correct’* trials and they would be indifferent to the variance of the distributions in ‘Ambiguous’ trials. We show that our monkeys deviate from such behaviours. The monkeys are more accurate on *‘Broad Correct’* trials than on *‘Narrow Correct’* trials (**Fig 3b, Supplementary Fig. 1**). Furthermore, in the *‘Ambiguous’* trials, the monkeys demonstrated a preference for the broadly distributed stream, which has greater variability across samples (**Fig 3c, Supplementary Fig. 1**). Such a pro-variance bias pattern of decision behaviour is similar to what was found in human subjects^19,20^ (**Fig 3d-e**).

**Figure 3.**
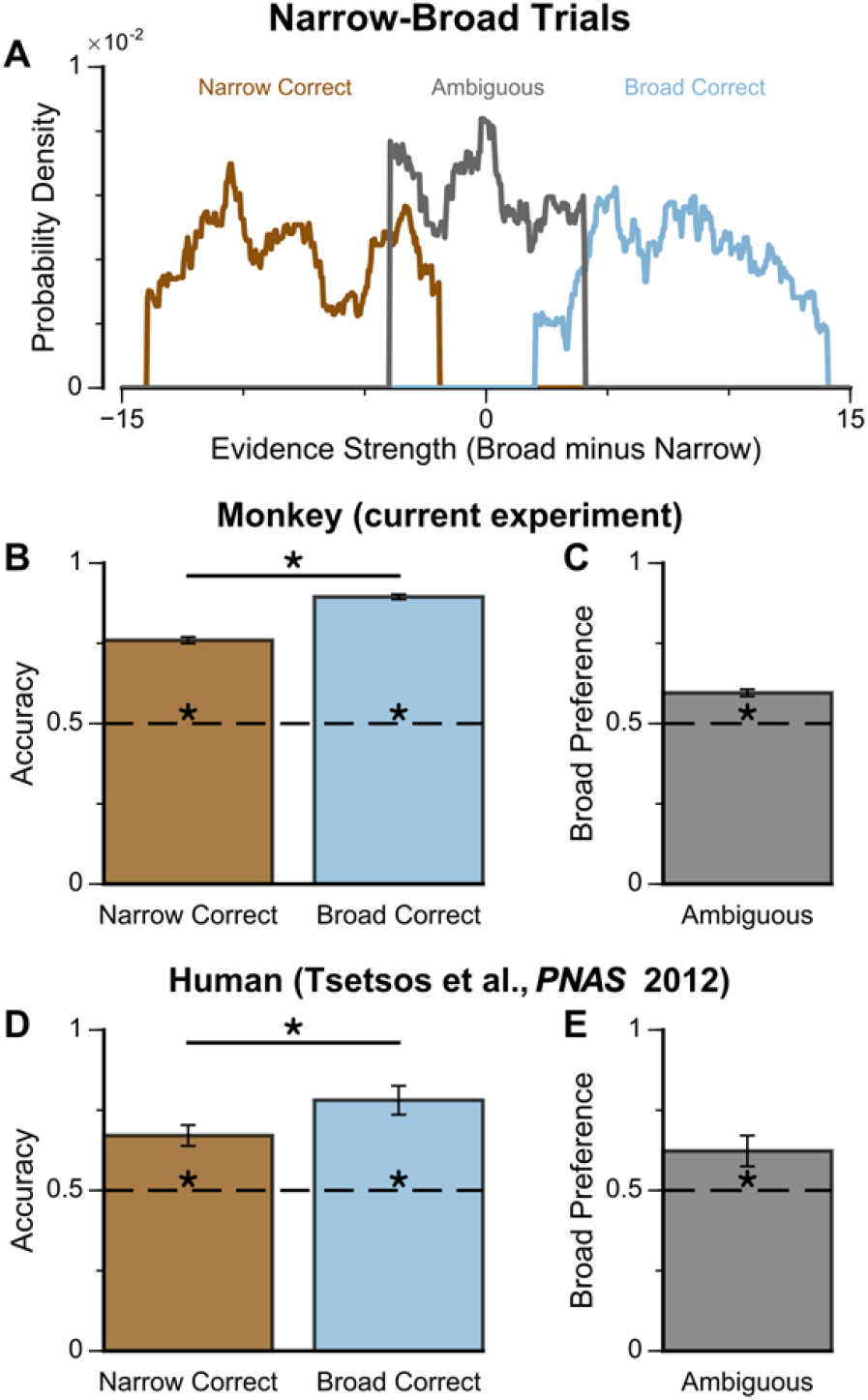
Subjects show a pro-variance bias in their choices on Narrow-Broad Trials, mirroring previous findings in human subjects. (**A**) The narrow-broad trials include three types of conditions, where either the narrow stream is correct (brown), the broad stream is correct (blue), or the difference in mean evidence is small (grey, ‘Ambiguous’ trials). See Methods and Supplementary Fig. 1 for details of the generating process. (**B-C**) Monkey choice performance on Narrow-Broad trials. (**B**) Subjects were significantly more accurate on ‘Broad-correct’ trials (Chi-squared test, chi = 99.05, p < 1×10^−10^). Errorbars indicate the standard error. (**C**) Preference for the broad option on ‘Ambiguous’ trials. Subjects were significantly more likely to choose the broad option (Binomial test, p < 1×10^−10^). (**D-E**) Human choice performance on Narrow-Broad trials previously reported by Tsetsos et al. 2012^20^. (**D**) Choice accuracy when either the narrow or the broad stream is correct, respectively. Subjects were more accurate on ‘Broad-correct’ trials. (**E**) Preference for the broad option on ‘Ambiguous’ trials. Subjects were more likely to choose the broad option.

To further probe the pro-variance bias, we studied choices from a larger pool of ‘Regular’ trials in which the mean evidences and variabilities of the two streams were set independently on each trial (**Fig4a, b, Supplementary Fig. 2**). ‘Regular’ trials allowed us to explore the pro-variance bias across a greater range of choice difficulties (**Fig 4c**) and quantitatively characterise its effect using regression analysis. On ‘Regular’ trials, subjects also demonstrated a preference for options with broadly distributed evidence. Regression analysis confirmed that evidence variability was a significant predictor of choice (**Fig 4d**; see **Methods**). In addition, we defined the pro-variance bias (PVB) index as the ratio of the regression coefficient for evidence standard deviation over the regression coefficient for mean evidence. This acted as a unitless measure of the pro-variance bias over the subjects’ sensitivity to the net evidence for choice selectivity. A PVB index value of 0 thereby indicates no pro-variance bias, whereas a PVB index value of 1 indicates the subject is as sensitive to evidence standard deviation as they are to mean evidence. The PVB index thus provides a quantitative measure of the pro-variance bias. From the ‘Regular’ trials, the PVB index across both monkeys was 0.173 (Monkey A = 0.230; Monkey H = 0.138).

**Figure 4.**
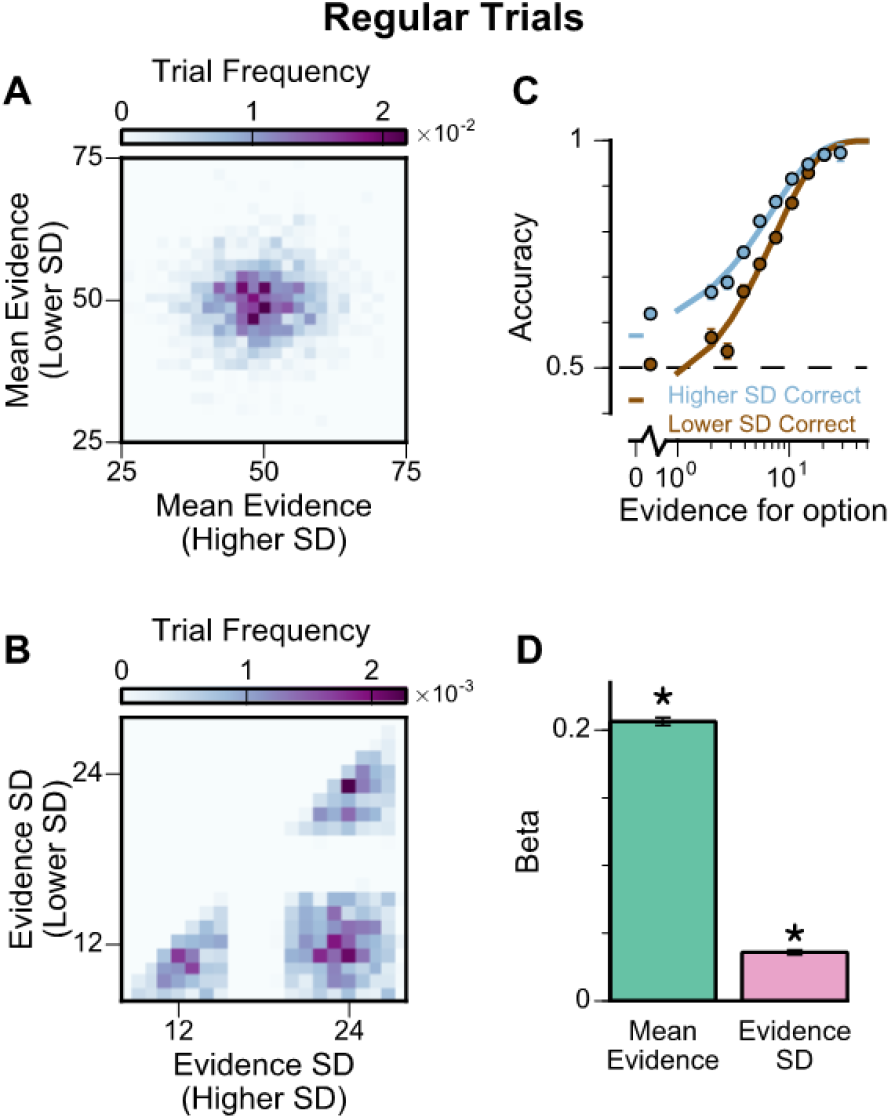
Subjects show a pro-variance bias in their choices on regular trials. For these analyses, stimulus streams were divided into ‘Lower SD’ or ‘Higher SD’ options post-hoc, on a trial-wise basis. (**A**) On regular trials, the mean evidence of each stream was independent. (**B**) Each stream is sampled from either a narrow or a broad distribution, such that about 50% of the trials have one broad stream and one narrow stream, 25% of the trials have two broad streams, and 25% of the trials have two narrow streams. (**C**) Psychometric function when either the ‘Lower SD’ (brown) or ‘Higher SD’ (blue) stream is correct in the regular trials. (**D**) Regression analysis using the left-right differences of the mean and standard deviation of the stimuli evidence to predict left choice. The beta coefficients quantify the contribution of both statistics to the decision-making processes of the monkeys (Mean Evidence: t = 74.78, p < 10^−10^; Evidence Standard Deviation: t = 19.65, p < 10^−10^). Notably, a significantly positive evidence SD coefficient indicates the subjects preferred to choose options which were more variable across samples. Errorbars indicate the standard error. For data separated by subjects, see **Supplementary Fig. 2**.

Recent work has suggested that when traditional evidence accumulation tasks are performed, it is hard to dissociate whether subjects are combining information across samples, or whether conventional analyses may be disguising a simpler heuristic^32,33^. In particular, an alternative decisionmaking strategy which does not involve temporal accumulation of evidence is to detect the single most extreme sample. Because the extreme sample will occur at different times in each trial, if a subject employed this strategy, the choice regression weights across time points would be distributed as in **Fig 2c,d**. Therefore, it is possible for these findings to be mistakenly interpreted as reflecting evidence accumulation. We wanted to quantitatively confirm that subjects were using the strategy we envisioned when designing our task, namely evidence accumulation. Additionally, we wanted to further investigate the relative contributions of mean evidence and evidence variability on choices. A logistic regression approach probed the influence upon choice of mean evidence, evidence variability, first/last samples, and the most extreme samples within each stream (**Supplementary Fig. 2e,h**, see **Methods**). A cross-validation approach revealed choice was principally driven by the mean evidence, verifying that subjects performed the task using evidence accumulation (**Supplementary Table 1**, see Methods).

Although this analysis revealed choices were not primarily driven by an ‘extreme sample detection’ decision strategy, another concern was whether partially employing this strategy could explain the pro-variance effect we observed. To address this, we compared the influence of ‘evidence variability’ versus the influence of ‘extreme samples’ on subjects’ choices. Cross-validation revealed that choices were better described by a model incorporating evidence variability, rather than the extreme sample values (**Supplementary Table 2**). We also demonstrated that including evidence variability as a co-regressor improved the performance of all combinations of nested models (**Supplementary Table 3**). In summary, it can be concluded that although subjects integrated across samples, they were additionally influenced by sample variability.

Existing algorithmic-level proposals for generating a pro-variance bias in human decision-making rely on the disregarding of sensory information before it enters the accumulation process, depending on its salience^19^. To investigate a possible alternative basis for the pro-variance bias, at the level of neural implementation, we sought to characterise decision-making behaviour in a biophysically-plausible spiking cortical circuit model (**Fig5a, b, Supplementary Fig. 3**)^21,34^. In the circuit architecture, two groups of excitatory pyramidal neurons are assigned to the left and right options, such that high activity in one group signals the response to the respective option. Excitatory neurons within each group are recurrently connected to each other via AMPA and NMDA receptors, and this recurrent excitation supports ramping activity and evidence accumulation. Both groups of excitatory neurons are jointly connected to a group of inhibitory interneurons, resulting in feedback inhibition and winner-take-all competition^21,22^. The two groups of excitatory neurons receive separate inputs-with each group receiving information about one of the two options (i.e. Group A receives IA reflecting the left option; Group B receives IB reflecting the right option). Specifically, we assume the bar heights from each stream are remapped, upstream of the simulated decision-making circuit, to evidence for the corresponding option depending on the cued context. Therefore, higher bars correspond to larger inputs in ‘ChooseHigh’ trials and smaller inputs in ‘ChooseLow’ trials. Combined together, this synaptic architecture endows the circuit model with decision-making functions.

**Figure 5:**
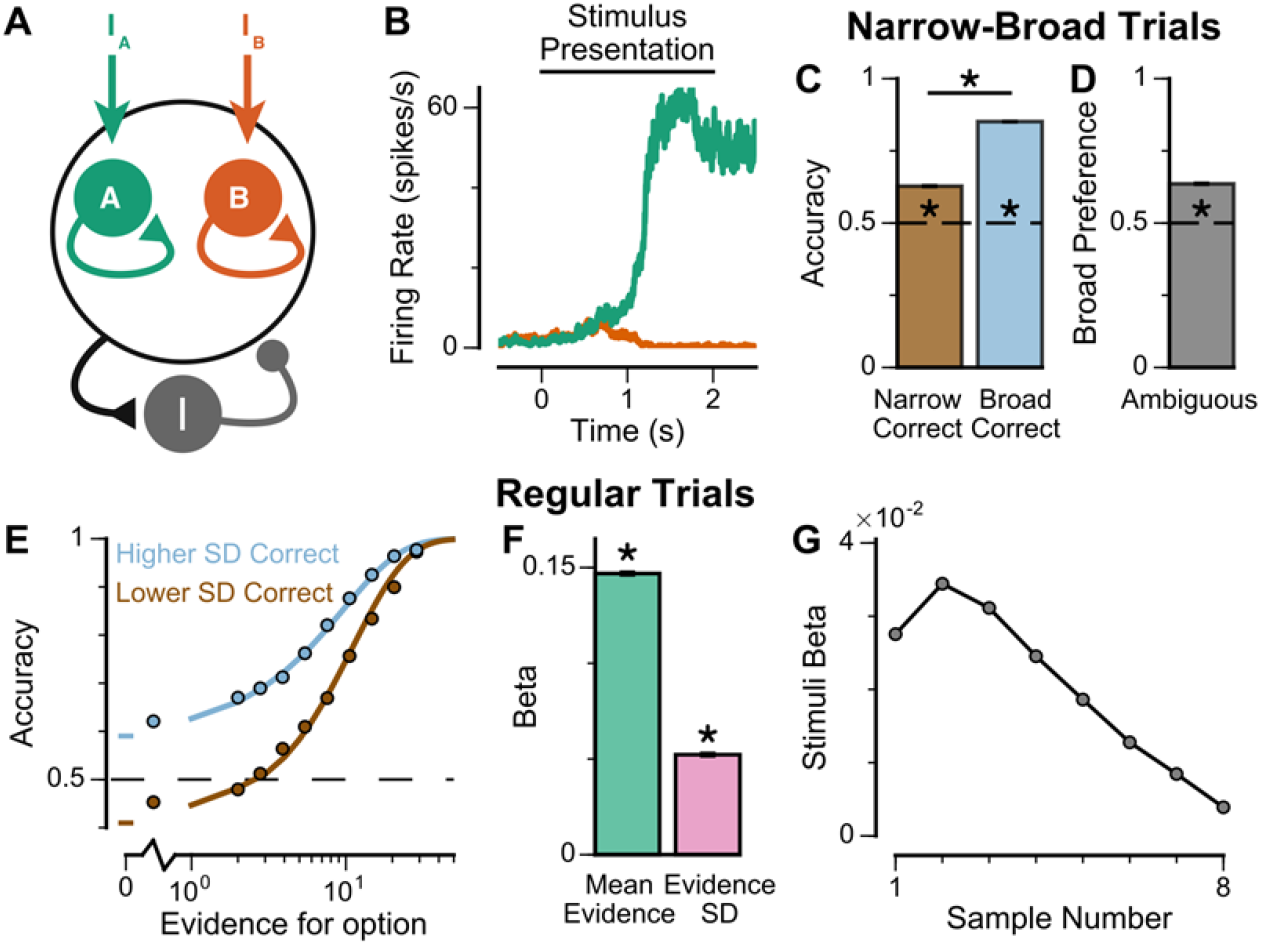
Spiking cortical circuit model reproduces pro-variance bias. (**A**) Circuit model schematic. The model consists of two excitatory neural populations which receive separate inputs (IA and IB), each reflecting the momentary evidence for one of the two stimuli streams. Each population integrates evidence due to recurrent excitation, and competes with the other via lateral inhibition mediated by a population of interneurons. (**B**) Example firing rate trajectories of the two populations on a single trial where option A is chosen. (**C, D**) Narrow-Broad Trials. (**C**) The circuit model is significantly more accurate when the broad stream is correct, than when the narrow stream is correct (Chi-squared test, chi = 1981, p < 1×10^−10^). (**D**) On ‘Ambiguous trials’, the circuit model is significantly more likely to choose the broad option (Binomial test, p < 1×10^−10^). (**E-G**) Regular trials. (**E**) The psychometric function of the circuit model when either the ‘Lower SD’ (brown) or ‘Higher SD’ (blue) stream is correct, respectively. (**F**) Regression analysis of the circuit model choices on regular trials, using evidence mean and variability as predictors of choice. Both quantities contribute to the decision-making process of the circuit model (Mean Evidence: t = 129.50, p < 10^−10^; Evidence Standard Deviation: t =45.27, p < 10^−10^). (**G**) Regression coefficients of the stimuli at different time-steps, showing the time course of evidence integration. The circuit demonstrates a temporal profile which decays over time, similar to the monkeys.

The spiking circuit model was tested with the same trial types as the monkey experiment. Importantly, not only can the circuit model perform the evidence accumulation task, it also demonstrated a provariance bias comparable to the monkeys (**Fig 5c-f**). Regression analysis showed that the circuit model utilises a strategy similar to the monkeys to solve the decision-making task (**Supplementary Fig. 3b**). The temporal process of evidence integration in the circuit model disproportionately weighted early stimuli over late stimuli (**Fig 5g**), similar to the evidence integration patterns observed in both monkeys. However, the circuit model demonstrated an initial ramp-up in stimuli weights due to the time needed for it to reach an integrative state.

To understand the origin of the pro-variance bias in the spiking circuit, we mathematically reduced the circuit model to a mean-field model (**Fig 6a**), which demonstrated similar decision-making behaviour to the spiking circuit (**Fig 6b, c, Supplementary Fig. 4**). The mean-field model, with two variables representing the integrated evidence for the two choices, allowed phase-plane analysis to further investigate the pro-variance bias. A simplified case was considered where the broad and narrow streams have the same mean evidence, and the stimuli evidence varies over time in the broad stream but not the narrow stream (i.e. σ_N_=0) (**Fig 6e-h**). This example provides an intuitive explanation for the pro-variance bias: a momentarily strong stimulus has an asymmetrically greater influence upon the decision-making process than a momentarily weak stimulus. It can be shown that such asymmetry arises from the expansive non-linearities of the firing rate profiles (**Fig 6d**).An advantage of the circuit model over existing algorithmic level explanations of the pro-variance bias is it can be used to make testable behavioural predictions in response to different synaptic or cellular perturbations, including E/I imbalance. In turn, perturbation experiments can constrain and refine model components. Therefore, we studied the behavioural effects of distinct E/I perturbations, and upstream sensory deficit, on decision making and in particular, pro-variance bias (**Fig 7, Supplementary Fig. 5**). Three perturbations were introduced to the circuit model: lowered E/I balance (via NMDA-R hypofunction on excitatory pyramidal neurons), elevated E/I balance (via NMDA-R hypofunction on inhibitory interneurons), or sensory deficit (as weakened scaling of external inputs to stimuli evidence) (**Fig 7a**).

**Figure 6:**
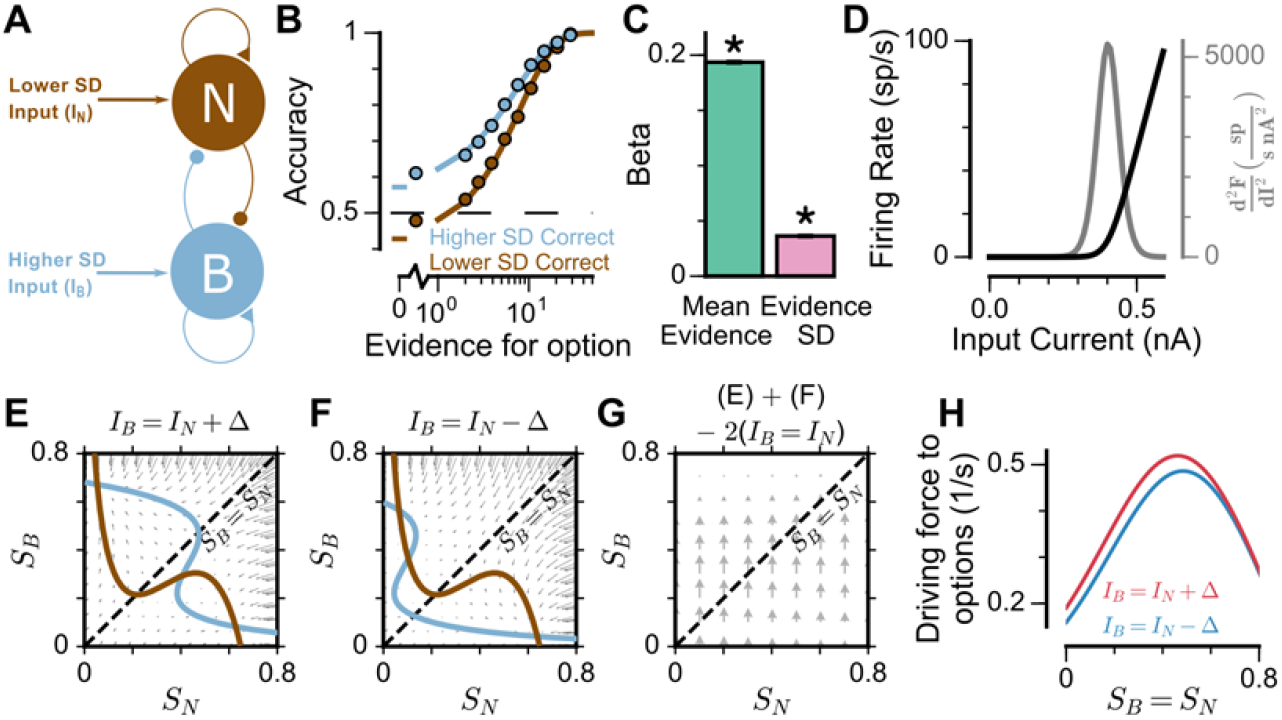
Mean-Field model explanation for pro-variance bias. (**A**) The mean-field model of the circuit, with two variables representing evidence for the two options. For simplicity, we assume one stream is narrow and one is broad, and label the populations receiving the inputs as N and B respectively. (**B**) Psychometric function of regular trials as in (**Fig 5E**). (**C**) Regression analysis of the regular trial data as in (**Fig 5F**) (mean: t = 143.42, p < 10^−10^; standard deviation: t =30.76, p < 10^−10^). (**D**) The mean-field model uses a generic firing rate profile (black), with zero firing rate at low inputs, then a near-linear response as input increases. Such profiles have an expansive non-linearity (with a positive second order derivative (grey)) that can generate pro-variance bias. (**E-H**) An explanation of the pro-variance bias using phase-plane analysis. (**E**) A momentarily strong stimulus from the broad stream will drive the model to choose broad (high S_B_, low S_N_). Blue and brown lines correspond to nullclines. (**F**) A momentarily weak stimulus in the broad stream will drive the model to choose narrow (high S_N_, low S_B_). (**G**) The net effect of one strong and one weak broad stimulus, compared with two average stimuli, is to drive the system to the broad choice. That is, a momentarily strong stimulus has an asymmetrically greater influence on the decision-making process than a momentarily weak stimulus, leading to pro-variance bias. (**H**) The net drive to the broad or narrow option when the broad stimulus is momentarily strong (red) or weak (blue), along the diagonal (S_B_ = S_N_ in **Fig 6G**).

**Figure 7:**
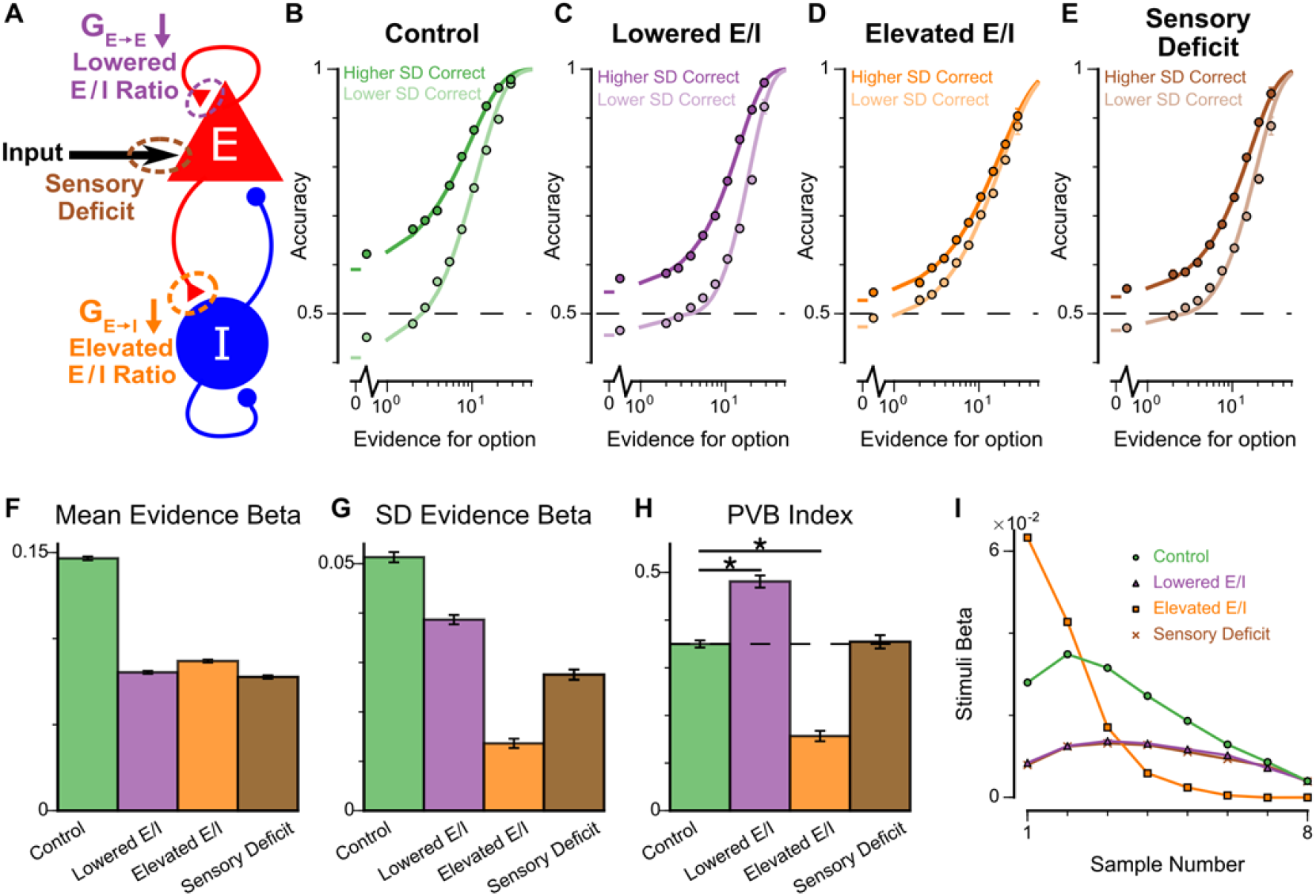
Predictions for E/I perturbations of the Spiking Circuit Model. (**A**) Model perturbation schematic. Three potential perturbations are considered: lowered E/I (via NMDA-R hypofunction on excitatory pyramidal neurons), elevated E/I (via NMDA-R hypofunction on inhibitory interneurons), or sensory deficit (as weakened scaling of external inputs to stimuli evidence).(**B-E**) The regular-trial choice accuracy for each of the circuit perturbations (dark colour for when the ‘Higher SD’ stream is correct, light colour for when the ‘Lower SD’ stream is correct).(**F-H**) Regression analysis on the regular trial choices of the four models, using evidence mean and evidence variability to predict choice.(**F**) The mean evidence regression coefficients in the four models. Lowering E/I, elevating E/I, and inducing sensory deficits similarly reduce the coefficient, reflecting a drop in choice accuracy. (**G**) The evidence standard deviation regression coefficients in the four models. All three perturbations reduce the coefficient, but to a different extent. (**H**) The PVB index (ratio of evidence standard deviation coefficient over mean evidence coefficient) provides dissociable predictions for the perturbations. The lowered E/I circuit increases the PVB index relative to the control model (permutation test, p <10^−5^), while the elevated E/I circuit decreases the PVB index (permutation test, p <10^−5^). The PVB index is roughly maintained in the sensory deficit circuit (permutation test, p = 0.695). The dashed line indicates the PVB index for the control circuit, * indicates p<10^−5^ when the PVB index is compared with the control circuit. (**I**) The regression weights of stimuli at different time-steps for the four models.

While all circuit models were capable of performing the task (**Fig 7b-e**), the choice accuracy of each perturbed model was reduced when compared to the control model. This was quantified by the regression coefficient of mean evidence (**Fig 7f**). In addition, the regression coefficient for evidence standard deviation was reduced for each perturbed model in comparison to the control model, indicating a lesser influence of evidence variability on choice (**Fig 7g**). Finally, in a dissociation between the three model perturbations, the PVB index was increased by lowered E/I, decreased by elevated E/I, and roughly unaltered by sensory deficits (**Fig 7h**). Further regression analyses indicated no obvious shift in utilised strategies relative to the control model (**Supplementary Fig. 5b**). In addition, the temporal weightings were distinctly altered by the elevated- and lowered-E/I perturbations (**Fig 7i**). The circuit model thus provided the basis of dissociable prediction by E/I-balance perturbing pharmacological agents.

To explore these predictions experimentally, we collected behavioural data from both monkeys following the administration of a subanaesthetic dose (0.5mg/kg, intramuscular injection) of the NMDA-R antagonist ketamine (see **Methods, Fig 8, Supplementary Fig. 6**). After a baseline period of the subjects performing the task, either ketamine or saline was injected intramuscularly (Monkey A: 13 saline sessions, 15 ketamine sessions; Monkey H: 17 saline sessions, 18 ketamine sessions). Administering ketamine had behavioural effects for around 30 minutes in both subjects. The data collected during this period formed a behavioural database of 4142 completed trials (Monkey A: 2276; Monkey H: 1866). Following ketamine administration, subjects’ choice accuracy was markedly decreased (**Fig 8a**), without a significant shift in their strategies (**Supplementary Fig. 6**, **Supplementary Table 4**). To understand the nature of this deficit, we studied the effect of drug administration on the pro-variance bias (**Fig 8b-f**). Although subjects were less accurate following ketamine injection, they retained a pro-variance bias (**Fig 8c**). Regression analysis confirmed ketamine caused choices to be substantially less driven by mean evidence (**Fig 8d**), but still strongly influenced by the standard deviation of evidence across samples (**Fig 8e**). The PVB index was significantly higher when ketamine was administered, than saline (permutation test p = 8×10^−6^, **Fig 8f**). Of all the circuit model perturbations, this was only consistent with lowered E/I balance (**Fig 7h**). Finally, we investigated the effect of ketamine on the time course of evidence weighting (**Fig 8g**). It caused a general downward shift of the temporal weights; but had no strong effects on how each stimulus was weighted relative to the others in the stream. This shifting of the weights could reflect a sensory deficit, but given the results of the pro-variance analysis, collectively the behavioural effects of ketamine are most consistent with a lowering of E/I balance.

**Figure 8:**
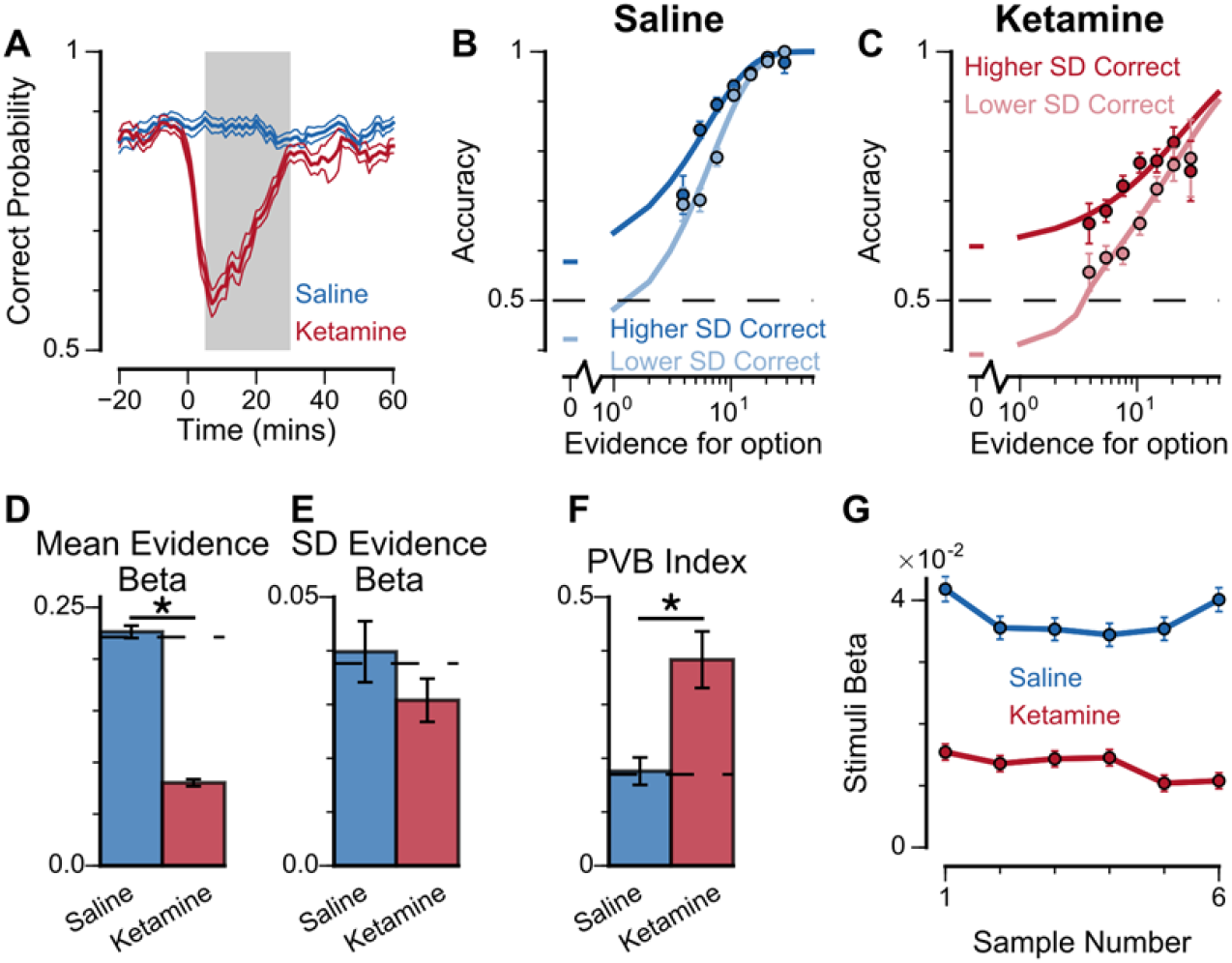
Experimental effects of ketamine on evidence accumulation behaviour produce an increased pro-variance bias, consistent with lowered excitation-inhibition balance. (**A**) Mean percentage of correct choices across sessions made by monkeys relative to the injection of ketamine (red) or saline (blue). Shaded region denotes ‘on-drug’ trials (trials 5-30 minutes after injection) which are used for analysis in the rest of the figure. (**B, C**) The psychometric function when either the ‘Lower SD’ or ‘Higher SD’ streams are correct, with saline (**B**) or ketamine (**C**) injection. (**D-F**) Ketamine injection impairs the decision-making of the monkeys, in a manner consistent with the prediction of the lowered E/I circuit model. Dashed lines indicate pre-injection values in each plot. (**D**) The regression coefficient for mean evidence, under injection of saline or ketamine. Ketamine significantly reduces the coefficient (permutation test, p < 1×10^−6^), reflecting a drop in choice accuracy. (E) The evidence standard deviation regression coefficient, under injection of saline or ketamine. Ketamine does not significantly reduce the coefficient (permutation test, p = 0.152). (**F**) Ketamine increases the PVB index (permutation test, p = 8×10^−6^), consistent with the model prediction of the lowered E/I circuit. (**G**) The regression weights of stimuli at different time-steps, for the monkeys with saline or ketamine injection. Ketamine injection lowers and flattens the curve of temporal weights, consistent with the lowered E/I circuit model. Errorbars in (**A**) indicate the standard error mean, in all other panels errorbars indicate the standard error. For data separated by subjects, see **Supplementary Fig. 6**.

## Discussion

Previous studies have shown human participants exhibit choice irrationalities when options differ in the standard deviation of the evidence samples, preferring choice options drawn from a more variable distribution^19,20^. By utilising a behavioural task with precise experimenter control over the distributions of time-varying evidence, we show that macaque monkeys exhibit a similar pro-variance irrationality in their choices akin to human participants. This pro-variance bias was also present in a spiking circuit model, which demonstrated a neural mechanism for this behaviour. We then introduced perturbations at distinct synaptic sites of the circuit, which revealed dissociable predictions for the effects of NMDA-R antagonism. Ketamine produced decision-making deficits consistent with a lowering of the cortical excitation-inhibition balance.

Biophysically grounded neural circuit modelling is a powerful tool to link cellular level observations to behaviour. Previous studies have shown recurrent cortical circuit models reproduce normative decision-making and working memory behaviour, and replicate the corresponding neurophysiological activity^21–26,35^. However, whether they are also capable of reproducing idiosyncratic cognitive biases has not previously been explored. Here we demonstrated pro-variance and primacy biases in a spiking circuit model. The primacy bias results from the formation of attractor states before all of the evidence has been presented. This neural implementation for bounded evidence accumulation corresponds with previous algorithmic explanations^29^.

The results from our spiking circuit modelling also provided a parsimonious explanation for the cause of the pro-variance bias within the evidence accumulation process. Specifically, strong evidence in favour of an option pushes the network towards an attractor state more so than symmetrically weak evidence pushes it away. In contrast, previous explanations for pro-variance bias proposed computations at the level of sensory processing upstream of evidence accumulation. In particular, a ‘selective integration’ model proposed that information for the momentarily weaker option is discarded before it enters the evidence accumulation process^19^. Crucially, our circuit model generated dissociable predictions for the effects of NMDA-R hypofunction on the pro-variance bias (PVB) index that were tested by follow-up ketamine experiments. While it is still unclear where and how in the brain the selective integration process takes place, our modelling results suggest that purely sensory deficits may not capture the alterations in choice behaviour observed under ketamine, in contrast to E/I perturbations in decision-making circuits (**Fig 7h**). Multiple complementary processes may simultaneously contribute to pro-variance bias during decision making, especially in complex behaviours over longer timescales. Future work will aim to contrast between these two models with neurophysiological data recorded while monkeys are performing this task.

Our pharmacological intervention experimentally verified the significance of NMDA-R function for decision-making. In the spiking circuit model, NMDA-Rs expressed on pyramidal cells are necessary for reverberatory excitation, without which evidence cannot be accumulated and stable working memory activity cannot be maintained. NMDA-Rs on interneurons are necessary for maintaining background inhibition and preventing the circuit from reaching an attractor state prematurely^21,25^. By administering ketamine, an NMDA-R antagonist, specific short-term deficits in choice behaviour were induced, which were consistent with a lowering of the cortical excitation-inhibition balance in the circuit model. This suggests the NMDA-R antagonist we administered systemically was primarily acting to inhibit neurotransmission onto pyramidal cells.

The physiological effects of NMDA-R antagonism on *in vivo* cortical circuits remains an unresolved question. A number of studies have proposed a net cortical disinhibition through NMDA-R hypofunction on inhibitory interneurons^4,10,11,36^. The disinhibition hypothesis is supported by studies finding NMDA-R antagonists mediate an increase in the firing of prefrontal cortical neurons, in rodents^37,38^ and monkeys^39–42^. On the other hand, the effects of NMDA-R antagonists on E/I balance may vary across neuronal sub-circuits within a brain area. For instance, in a working memory task, ketamine was found to increase spiking activity of response-selective cells, but decrease activity of the task-relevant delay-tuned cells in primate prefrontal cortex^26^. Such specificity might explain why several studies reported less conclusive effects of NMDA-R antagonists on overall prefrontal firing rates in monkeys^26,43^. *In vitro* work has also revealed the excitatory post-synaptic potentials (EPSPs) of prefrontal pyramidal neurons are much more reliant on NMDA-R conductance than parvalbumin interneurons^44^. Other investigators combining neurophysiological recordings with modelling approaches have also concluded that the action of NMDA-R antagonists is primarily upon pyramidal cells^26,45^. Our present findings, integrating pharmacological manipulation of behaviour with biophysically-based spiking circuit modelling, suggest that the ketamine-induced behavioural biases are more consistent with a lowering of excitation-inhibition balance. Future work with electrophysiological recordings during the performance of our task, under pharmacological interventions, can potentially dissociate the effect of ketamine on E/I balance specifically in cortical neurons exhibiting decision-related signals.

The minutes-long timescale of the NMDA-R mediated decision-making deficit we observed was also consistent with the psychotomimetic effects of subanaesthetic doses of ketamine in healthy humans^7,11^. As NMDA-R hypofunction is hypothesised to play a role in the pathophysiology of schizophrenia^5,6,10,11^, our findings have important clinical relevance. Previous studies have demonstrated impaired perceptual discrimination in patients with schizophrenia performing the random-dot motion (RDM) decision-making task^15–17^. Although the RDM has predominantly been used to study evidence accumulation^13^, previously this performance deficit in schizophrenia was interpreted as reflecting a diminished representation of sensory evidence in visual cortex^15,18^. Based on our task with precise temporal control of the stimuli, our findings suggest that NMDA-R antagonism alters the decision-making process in association cortical circuits. Dysfunction in these association circuits may therefore provide an important contribution to cognitive deficits – one that is potentially complementary to upstream sensory impairment. Crucially, our task uniquely allowed us to rigorously verify that the subjects used an accumulation strategy to guide their choices (cf. previous animal studies^13,14,46–48^), with these analyses suggesting the strategy our subjects employed was consistent with findings in human participants. This consistency further ensures our findings may translate across species, in particular to clinical populations.

Another related line of schizophrenia research has shown a decision-making bias known as jumping to conclusions (JTC)^49,50^. The JTC has predominately been demonstrated in the ‘beads task’, a paradigm where participants are shown two jars of beads, one mostly pink and the other mostly green (typically 85%). The jars are hidden, and the participants are presented a sequence of beads drawn from a single jar. Following each draw, they are asked if they are ready to commit to a decision about which jar the beads are being drawn from. Patients with schizophrenia typically make decisions based on fewer beads than controls. Importantly, this JTC bias has been proposed as a mechanism for delusion formation. Based on the JTC literature, one plausible hypothesis for behavioural alteration under NMDA-R antagonism in our task may be a strong increase in the primacy bias, whereby only the initially presented bar samples would be used to guide the subjects’ decisions. However, following ketamine administration, we did not observe a strong primacy – instead all samples received roughly the same weighting. There are important differences between our task and the beads task. In our task, the stimulus presentation is shorter (2 seconds, compared to slower sampling across bead draws), and is of fixed duration rather than terminated by the subject’s choice, and therefore may not involve the perceived sampling cost of the beads task^51^.

Our precise experimental paradigm and complementary modelling approach allowed us to meticulously quantify how monkeys weight time-varying evidence and robustly dissociate sensory and decision-making deficits – unlike prior studies using the RDM and beads tasks. Our approach can be readily applied to experimental and clinical studies to yield insights into the nature of cognitive deficits and their potential underlying E/I alterations in pharmacological manipulations and pathophysiologies across neuropsychiatric disorders, such as schizophrenia^52,53^ and autism^52,54–56^. Finally, our study highlights how precise task design, combined with computational modelling, can yield translational insights across species, including through pharmacological perturbations, and across levels of analysis, from synapses to cognition.

## Methods

### Subjects

Two adult male rhesus monkeys (*M. mulatta*), subjects A and H, were used. The subjects weighed 12–13.3 kg, and both were ~6 years old at the start of the data collection period. We regulated their daily fluid intake to maintain motivation in the task. All experimental procedures were approved by the UCL Local Ethical Procedures Committee and the UK Home Office, and carried out in accordance with the UK Animals (Scientific Procedures) Act.

### Behavioural protocol

Subjects sat head restrained in a primate behavioural chair facing a 19-inch computer screen (1,280 × 1024-px screen resolution, and 60-Hz refresh rate) in a dark room. The monitor was positioned 59.5 cm away from their eyes, with the height set so that the centre of the screen aligned with neutral eye level for the subject. Eye position was tracked using an infrared camera (ISCAN ETL-200) sampled at 240 Hz. The behavioural paradigm was run in the MATLAB-based toolbox MonkeyLogic (http://www.monkeylogic.net/, Brown University)^57–59^. Eye position data was relayed to MonkeyLogic for use online during the task, and was recorded for subsequent offline analysis. Following successful trials, juice reward was delivered to the subject using a precision peristaltic pump (ISMATEC IPC). Subjects performed two types of behavioural sessions: standard and pharmacological. In pharmacological sessions, following a baseline period, either an NMDA-R antagonist (Ketamine) or saline was administered via intramuscular injection. Monkey A completed 41 standard sessions, and 28 pharmacological sessions (15 ketamine; 13 saline). Monkey H completed 68 standard sessions, and 35 pharmacological sessions (18 ketamine; 17 saline).

### Injection protocol

Typically, two pharmacological sessions were performed each week, at least 3 days apart. Subjects received either a saline or ketamine injection into the trapezius muscle while seated in the primate chair. Approximately 12 minutes into the session, local anaesthetic cream was applied to the muscle. At 28 minutes, the injection was administered. The task was briefly paused for this intervention (64.82 +/− 10.85 secs). Drug dose was determined through extensive piloting, and a review of the relevant literature^26,60^. The dose used was 0.5mg/kg.

### Task

Subjects were trained to perform a two-alternative value-based decision-making task. A series of bars, each with different heights, were presented on the left and right-side of the computer monitor. Following a post-stimulus delay, subjects were rewarded for saccading towards the side with either the higher or lower average bar-height, depending upon a contextual cue displayed at the start of the trial (see **Fig 1a** inset). The number of pairs of bars in each series was either four (*‘ShortSampleTrial’*) or eight (*‘LongSampleTrial’*) during trials in each standard behavioural session. In this report, we only consider the results from the eight sample trials, though similar results were obtained from the four sample trials. The number of bars was always six during pharmacological sessions.

The bars were presented inside of fixed-height rectangular placeholders (width, 84px; height, 318px). The placeholders had a black border (thickness 9px), and a grey centre where the stimuli were presented (width, 66px; height, 300px). The bar heights could take discrete percentiles, occupying between 1% and 99% of the grey space. The height of the bar was indicated by a horizontal black line (thickness 6px). Beneath the black line, there was 45° grey gabor shading.

An overview of the trial timings is outlined in **Fig 1a**. Subjects initiated a trial by maintaining their gaze on a central, red fixation point for 750ms. After this fixation was completed, one of four contextual cues (see **Fig 1a** inset) was centrally presented for 350ms. Subjects had previously learned that two of these cues instructed to choose the side with the higher average bar-height (*‘ChooseHighTrial’*), and the other two instructed to choose the side with the lower average bar-height (*‘ChooseLowTrial’*). Next, two black masks (width, 84px; height, 318px) were presented for 200ms in the location of the forthcoming bar stimuli. These were positioned either side of the fixation spot (6° visual angle from centre). Each bar stimulus was presented for 200ms, followed by a 50ms inter-stimulus-interval where only the fixation point remained on the screen. Once all of the bar stimuli had been presented, the mask stimuli returned for a further 200ms. There was then a post stimulus delay (250-750ms, uniformly sampled across trials). Following this, the colour of the fixation point was changed to green (go cue), and two circular saccade targets appeared on each side of the screen where the bars had previously been presented. This cued the subject to indicate their choice by making a saccade to one of the targets. Once the subject reported their decision, there were two stages of feedback. Immediately following choice, the green go cue was extinguished, the contextual cue was re-presented centrally, along with the average bar heights of the two series of stimuli previously presented. The option the subject chose was indicated by a purple outline surrounding the relevant bar placeholder (width, 3.8°; height, 10°). Following 500ms, the second stage of feedback began. The correct answer was indicated by a white outline surrounding the bar placeholder (width, 5.7°; height, 15°). On correct trials, the subject was rewarded for a length of time proportional to the average height of the chosen option (directly proportional on a *‘ChooseHighTrial’*, negatively proportional on a *‘ChooseLowTria?*). On incorrect trials, there was no reward. Regardless of the reward amount, the second feedback stage lasted 1200ms. This was followed by an inter-trial-interval (1.946+/− 0.051 secs; for Standard Session, across all completed included trials). The inter-trial-interval duration was longer on *‘ShortSampleTrials’* than ‘LongSampleTrials’, in order for the trials to be an equal duration, and facilitate a similar reward rate between the two conditions.

Subjects were required to maintain central fixation from the fixation period until they indicated their choice. If the initial fixation period was not completed, or fixation was subsequently broken, the trial was aborted and the subject received a 3000ms timeout (Trials in standard sessions: Monkey A – 22.46%, Monkey H – 15.27%). On the following trial, the experimental condition was not repeated. If subjects failed to indicate their choice within 8000ms, a 5000ms timeout was initiated (Trials in standard sessions: Monkey A – 0%, Monkey H – 0%).

Experimental conditions were blocked according to the contextual cue and evidence length. This produced four block types (*ChooseHighShortSampleTrial (H4), ChooseHighLongSampleTrial (H8), ChooseLowShortSampleTrial (L4), ChooseLowLongSampleTrial (L8*)). At the start of each session, subjects performed a short block of memory-guided saccades (MGS)^61^, completing 10 trials. Data from these trials is not presented in this report. Following the MGS block, the first block of decision-making trials was selected at random. After the subject completed 15 trials in a block, a new block was selected without replacement. Each new block had to have either the same evidence length or the same contextual cue as the previous block. After all four blocks had been completed, there was another interval of MGS trials. A new evidence accumulation start block was then randomly selected. As there were four block types, and either the evidence length or the contextual cue had to be preserved across a block switch, there were two ‘sequences’ in which the blocks could transition (i.e. H4→H8→L8→L4; or H4→L4→L8→H8, if starting from H4). Following the intervening MGS trials, the blocks transitioned in the opposite sequence to those used previously, starting from the new randomly chosen block. This block switching protocol was continued throughout the session. At the start of each block, the background of the screen was changed for 5000ms to indicate the evidence length of the forthcoming block. A burgundy colour indicated an 8 sample block was beginning, a teal colour indicated a 4 sample block was beginning.

### Trial Generation

The heights of the bars on each trial were precisely controlled. On the majority of trials (Regular Trials, Completed trials in standard sessions: Monkey A – 76.67%, Monkey H – 76.23%), the heights of each option were generated from independent Gaussian distributions (**Fig 4a, b**). There were two levels of variance for the distributions, designated as ‘Narrow’ and ‘Broad’. The mean of each distribution, μ, was calculated as μ = 50 + Z*σ, where 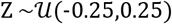, and σ was either 12 or 24 for narrow and broad stimuli streams. The individual bar heights were then determined by 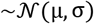. The trial generation process was constrained so the samples reasonably reflected the generative parameters. These restrictions required bar heights to range from 1 to 99, and the actual σ for each stream to be no more than 4 from the generative value. On any given trial, subjects could be presented with two narrow streams, two broad streams, or one of each. The evidence variability was therefore independent between the two streams. For post-hoc analysis (**Fig 4**) we defined one stream as the ‘Lower SD’ option on each trial, and the other the ‘Higher SD’ option, based upon the sampled/actual σ.

A proportion of ‘irrationality trials’ were also specifically designed to elucidate the effects of evidence variability on choice, and whether subjects displayed primacy/recency biases^20^. These trials occurred in equal proportions within all four block types. Only one of these irrationality trial types was tested in each behavioural session.

Narrow-broad trials (Completed trials in standard sessions: Monkey A – 14.87%, Monkey H – 15.78%) probed the effect of evidence variability on choice^20^. Within this category of trials, there were three conditions (**Fig 3a**). In each, the bar heights of one alternative were associated with a narrow Gaussian distribution 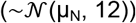, and the bar heights from the other with a broad Gaussian distribution 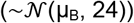. In the first two conditions, *‘Narrow Correct’* 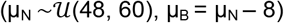 and *‘Broad Correct’* 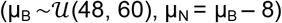, there was a clear correct answer. In the third condition, *‘Ambiguous’* 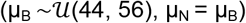, there was only small evidence in favour of the correct answer. In all of these conditions, the generated samples had to be within 4 of the generating σ. Furthermore, on ‘Narrow Correct’ and ‘Broad Correct’ trials the difference between the mean evidence of the intended correct and incorrect stream had to range from +2 to +14. On the ‘Ambiguous’ trials, the mean evidence in favour of one option over the other was constrained to be <4. A visualisation of the net evidence in each of these trial types is displayed (**Fig 3a**). For the purposes of illustration, the probability density was smoothed by a sliding window of ±1, within the generating constraints described above (‘Narrow Correct’ and ‘Broad Correct’ trials have net evidence for correct option within [2, 14]; ‘Ambiguous’ trials have net evidence within [-4, 4]). A very small number of trials were excluded from this visualisation, because their net evidence fell marginally outside the constraints. This was because bar heights were rounded to the nearest integer (due to the limited number of pixels on the computer monitor) after the generating procedure and the plot reflects the presented bar heights.

Half-half trials (Completed trials in standard sessions: Monkey A – 8.46%, Monkey H – 8.00%) probed the effect of temporal weighting biases on choice^20^. The heights of each option were generated using the same Gaussian distribution 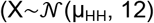, where 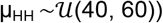. This distribution was truncated to form two distributions: X_High_ {mean(X)– 0.5*SD(X),∞}, and X_Low_ {-∞, mean(X)+ 0.5*SD(X)}. On each trial, one option was designated *‘HighFirst’* – where the first half of bar heights was drawn from X_High_ and the second half of bar heights drawn from X_Low_. This process was also constrained so that the mean of samples drawn from X_High_ had to be at least 7.5 greater than those taken from X_Low_. The other option was *‘LowFirst’*, where the samples were drawn from the two distributions in the reverse order.

### Task Modifications for Pharmacological Sessions

Minor adjustments were made to the task during the pharmacological sessions to maximise trial counts available for statistical analysis. Trial length was fixed to 6 pairs of samples. The block was switched between *‘ChooseHigh6Sample’* and *‘ChooseLow6Sample’* after 30 completed trials, without intervening MGS trials. From our pilot data, it was clear ketamine reduced choice accuracy. In order to maintain subject motivation, the most difficult ‘Regular’ and ‘HalfHalf’ trials were not presented. Following the trial generation procedures described above, in pharmacological sessions these trials were additionally required to have >4 mean difference in evidence strength. Of the *‘Narrow-Broad’* trials, only ‘*Ambiguous’* conditions were used; but no further constraints were applied to these trials. In some sessions, a small number of control trials were used, in which the bar heights for each option were fixed across all of the samples. All analyses utilised ‘Regular’, ‘Half-Half’, and ‘Narrow-Broad’ trials. Monkey H did not always complete sufficient trials once ketamine was administered. Sessions where the number of completed trials was fewer than the minimum recorded in the saline sessions were discarded (6 of 18 sessions). Following ketamine administration, Monkey A did not complete fewer trials in any session than the minimum recorded in a saline session.

### Behavioural Data Analysis

To assess decision-making accuracy during standard sessions, we initially fitted a psychometric function^14,29^ to subjects’ choices pooled across ‘Regular’ and ‘Narrow-Broad’ trials (**Fig 2a, b**). This defines the choice accuracy (*P*) as a function of the difference in mean evidence in favour of the correct choice (evidence strength, *x*):

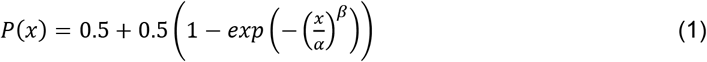

where *α* and *β* are respectively the discrimination threshold and order of the psychometric function, and *exp* is the exponential function. To illustrate the effect of pro-variance bias, we also fitted a three-parameter psychometric function to the subjects’ probability to choose the higher SD option (*P_HSD_*) in the ‘Regular’ trials, as a function of the difference in mean evidence in favour of the higher SD option on each trial (*x_HSD_*):

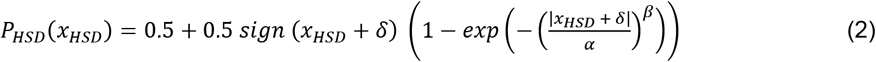

where *δ* is the psychometric function shift, and *sign* returns 1 and −1 for positive and negative inputs respectively.

In both cases, the psychometric function is fitted using the method of maximum-likelihood estimation (MLE), with the estimator

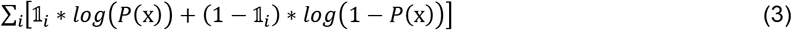

(and similarly for *P_HSD_* & *x_HSD_*), where *i* is summed across trials. 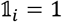 if the correct (higher SD) option is chosen in trial *i* and 0 otherwise.

The temporal weights of stimuli were calculated using logistic regression. This function defined the probability (PL) of choosing the left option:

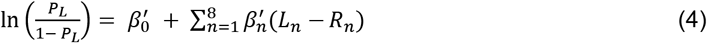

where 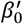 is a bias term, 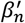 reflects the weighting given to the nth pair of stimuli, *L_n_* and *R_n_* reflect the evidence for the left and right option at each time point.

Regression analysis was used to probe the influence of evidence mean, and evidence variability on choice during the ‘Regular’ trials (**Fig 4d, 5f, 6c, 7f-h, 8d-f, Supp 2d,g, Supp 6c,h**). This function defined the probability (P_L_) of choosing the left option:

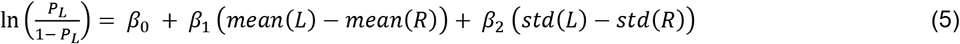

where *β*_0_ is a bias term, *β*_1_ reflects the influence of evidence mean, and *β*_2_ reflects the influence of standard deviation of evidence (evidence variability).

This approach was extended to probe other potential influences on the decision-making process. An expanded regression model was defined as follows:

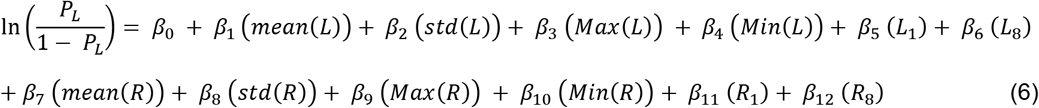

where *β*_0_ is a bias term, *β*_1_ reflects the influence of evidence mean of the left samples, *β*_2_ reflects the influence of evidence variability of the left samples, *β*_2_ reflects the influence of the maximum left sample, *β*_4_ reflects the influence of the minimum left sample, *β*_5_ reflects the influence of the first left sample, *β_6_* reflects the influence of the last left sample. *β*_7_ to *β*_12_ reflect the same attributes for samples on the right side of the screen. Due to strong correlations among evidence standard deviation, maximum, and minimum, the regression model without *β*_2_ and *β*_8_ is used to evaluate the contribution of regressors other than evidence mean and standard deviation to the decision making process (**Fig Supp 2e,h, Supp 3b, Supp 4b, Supp 5b, Supp 6d,i**).

The goodness-of-fit of various regression models with combinations of the predictors in the full model (equation 6) were compared using a 10-fold cross-validation procedure (**Supplementary Tables 1-4**). Trials were initially divided into 10 groups. Data from 9 of the groups was used to train each regression model and calculate regression coefficients. The likelihood of the subjects’ choices in the left-out group (testing group), given the regression coefficients, could then be determined. The loglikelihood was then summed across these left-out trials. This process was repeated so that each of the 10 groups acted as the testing group. The whole cross-validation procedure was performed 100 times, and the average log-likelihood values were taken.

To initially explore the time course of drug effects on decision-making, we plotted choice accuracy (combined across ‘Regular’, ‘Half-Half’ and ‘Narrow-Broad’ trials) relative to drug administration (**Fig 8a**). Trials were binned relative to the time of injection. Within each session, choice accuracy was estimated at every minute, using a 6-minute window around the bin centre. Accuracy was then averaged across sessions. To further probe the influence of drug administration on decision-making, we defined an analysis window based upon the time course of behavioural effects. All trials before the time of injection were classified as ‘pre-drug’. All trials beginning 5-30 minutes after injection were defined as ‘on-drug’ trials. These trials were then analysed using the same methods as described for the Standard sessions.

To quantify the effect of ketamine administration on the PVB index (**Fig 8f, Fig Supp 6c,h**), we performed a permutation test. Trials collected during ketamine administration were compared with those collected during saline administration. The test statistic was calculated as the difference between the PVB index in ketamine and saline conditions. For each permutation, trials from the two sets of data were pooled together, before two shuffled sets with the same number of trials as the original ketamine and saline data were extracted. Next, the PVB index was computed in each permuted set, and the difference between the two PVB indices calculated. The difference measure for each permutation was used to build a null distribution with 1000000 entries. The difference measure from the true data was compared with the null distribution to calculate a p-value.

### Spiking Circuit Model

A biophysically-based spiking circuit model was used to replicate decision making dynamics in a local association cortical microcircuit. The model was based on^21^, but with minor modifications from a previous study^34^. The current model had one extra change in the input representation of the stimulus, described in detail below.

The circuit model consisted of *N_E_* = 1600 excitatory pyramidal neurons and *N_I_* = 400 inhibitory interneurons, all simulated as leaky integrate-and-fire neurons. All neurons were recurrently connected to each other, with NMDA and AMPA conductances mediating excitatory connections, and GABAA conductances mediating inhibitory connections. All neurons also received background inputs, while selective groups of excitatory neurons (see below) received stimulus inputs. Both background and stimulus inputs were mediated by AMPA conductances with Poisson spike trains.

Within the population of excitatory neurons were two non-overlapping groups of size *N_E,G_* = 240. Neurons within the two groups received separate inputs reflecting the left and right stimuli streams. Neurons in the same group preferentially connected to each other (with a multiplicative factor *w*_+_ > 1 to the connection strength), allowing integration of the stimulus input. The connection strength to any other excitatory neurons was reduced by a factor *w*_−_ < 1 in a manner which preserved the total connection strength. Due to lateral inhibition mediated by interneurons, excitatory neurons in the two different groups competed with each other. Inhibitory neurons, as well as excitatory neurons not in the two groups, were insensitive to the presented stimuli and were non-selective toward either choices or the respective neuron groups.

Momentary stimuli bar evidences were modelled as Poisson inputs (from an upstream sensory area) to the two groups of excitatory neurons (**Fig 5a**). The mean rate of Poisson input for any group, *μ*, linearly scaled with the corresponding stimulus evidence:

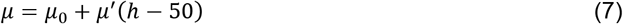

where *h* ∈ [0,100] represented the momentary stimulus evidence, equal to the bar height in ChooseHigh trials, and 100 minus bar height in ChooseLow trials. *μ*_0_ = 30*Hz* was the input strength when h = 50, and *μ*’ = 1*Hz*. For simplicity, we assumed each bar stimulus lasted 250ms, rather than 200ms with a subsequent 50ms inter-stimuli interval as in the experiment.

The circuit model simulation outputs spike data for the two excitatory populations, which are then converted to population activity smoothened with a 0.001s time-step via a casual exponential filter. In particular, for each spike of a given neuron, the histogram-bins corresponding to times before that spike receives no weight, while the histogram-bins corresponding to times after the spike receives a weight of 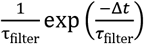, where Δ*t* is the time of the histogram-bin after the spike, and *τ*_filter_=20ms.

From the population activity of the two excitatory populations, a choice is selected 2s after stimulus offset, based on the population with higher activity. Stimulus inputs in general drive categorical, winner-take-all competitions such that the winning population will ramp up its activity until a high attractor state (>30Hz, in comparison to approximately 1.5Hz baseline firing rate), while suppressing the activity of the other population below baseline via lateral inhibition (**Fig 5b**). It is also possible that neither population reaches the high-activity state. Both populations, remaining at the spontaneous state, will have similarly low activities, such that the decision readout is random.

In addition to the control model, three perturbed spiking circuit models were considered^25,34^: lowered E/I balance, elevated E/I balance, and sensory deficit. E/I perturbations were implemented through hypofunction of NDMARs (**Fig 7a**), as this is a leading hypothesis in the pathophysiology of schizophrenia^4,5,10^. NMDA-R antagonists such as ketamine also provide a leading pharmacological model of schizophrenia^7,11^. NMDA-R hypofunction on excitatory neurons (reduced *G_E→E_*) resulted in lowered E/I ratio, whereas NMDA-R hypofunction on interneurons (reduced *G_E→I_*,) resulted in elevated E/I ratio due to disinhibition^34^. Sensory deficit was implemented as weakened scaling of external inputs to stimuli evidence, resulting in reduced *μ*. For the exact parameters, the lowered E/I model reduced *G_E→E_* by 1.75%, the elevated E/I model reduced *G_E→I_* by 3.5%, and the sensory deficit model had *μ*’ = 0.74*Hz*.

Each of the four circuit models completed 94,000 ‘Regular’ trials, where both streams are narrow in 25% of the trials, both streams are broad in 25% of the trials, and one stream is narrow and one is broad in 50% of the trials. All trials were generated identically as in standard session experiments. The control model also completed 47,000 standard session Narrow-Broad trials. The same permutation test described earlier for comparing PVB index between ketamine and saline conditions was also used to quantify whether various perturbed circuit models have different PVB indices relative to the control model (**Fig 7h**).

### Mean Field Model

The current spiking circuit model was mathematically reduced to a mean-field model, as outlined in^62^, in the same manner as from^21^ to^22^. The mean-field model consisted of two variables, namely the NMDA-R gating variables of the two groups of excitatory neurons, which represented the integrated evidence for the two choices. Using phase-plane analysis, the mean-field model provided an intuitive explanation for the pro-variance bias (see **Fig 6**).

The mean-field model completed 94,000 standard session ‘Regular’ trials, in the same manner as the circuit models.

### Code and Data Availability

Stimuli generation and data analysis for the experiment were performed in MATLAB. The spiking circuit model was implemented using the Python-based Brian2 neural simulator^63^, with a simulations time step of 0.02ms. Further analyses for both experimental and model data were completed using custom-written Python and MATLAB codes. All codes are available from the authors upon reasonable request.

## Supplementary Figures

**Supplementary Figure 1.**
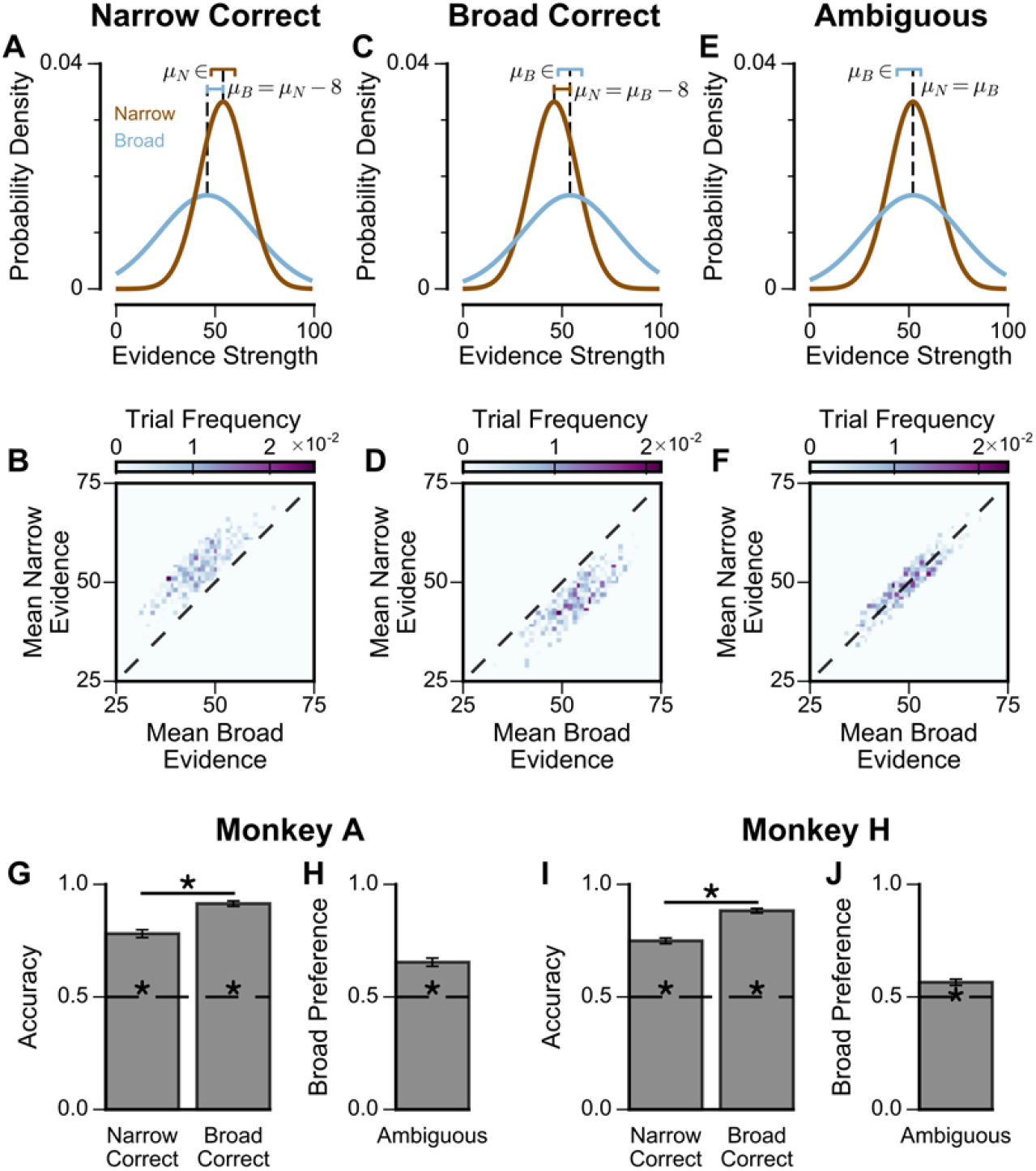
Extra Information on Narrow-Broad Trials: (**A**) The generating process of the narrow-correct trials, for each narrow (brown) and broad (blue) stimuli sample. A full stream sequentially presents 8 such stimuli, each for 200ms with a 50ms inter-sample interval in between. In each trial where the narrow choice is correct, the generating mean of the narrow stream, μ_N_, is uniformly sampled from [48,60]. The generating mean of the broad stream, μ_B_, is then set to be μ_N_ – 8. For all trials, the generating standard deviation of the narrow and broad streams are σ_N_ = 12, σ_B_ = 24 respectively. The lines above the distributions denote the ranges of μ_N_ and μ_B_. The particular values of μ_N_ and μ_B_ in this figure are shown for one trial, and chosen arbitrarily for illustration purpose. Given the generating means and standard deviations in a trial, a sequence of 8 stimuli samples are generated from a Gaussian process with certain constraints, for each of the narrow and broad options (See **Methods**). (**B**) Sampled distribution of the mean evidence of the narrow and broad streams, across all trials for both monkeys where the narrow option is correct. (**C, D**) Same as (**A, B**) but for broad-correct trials. Here, μ_B_ is uniformly sampled from [48,60], and μ_N_ is set to be μ_B_ −8. (**E, F**) Same as (**A, B**) but for ambiguous trials. Here, μ_N_and μ_B_ are equal and uniformly sampled from [44, 56]. (**G**) The accuracy of Monkey A in the narrow-correct and broad-correct trials. Monkey A was significantly more accurate on ‘Broad-correct’ trials (Chi-squared test, chi = 38.39, p = 5.80×10^−10^). Errorbars show the standard error. (**H**) The probability for Monkey A to choose the broad option in ambiguous trials. Monkey A was significantly more likely to choose the broad option (Binomial test, p < 1×10^−10^). (**I**) Same as (**G**) but for Monkey H (Chi-squared test, chi = 59.46, p < 1×10^−10^). (**J**) Same as (**H**) but for Monkey H (Binomial test, p = 3.00×10^−6^).

**Supplementary Figure 2.**
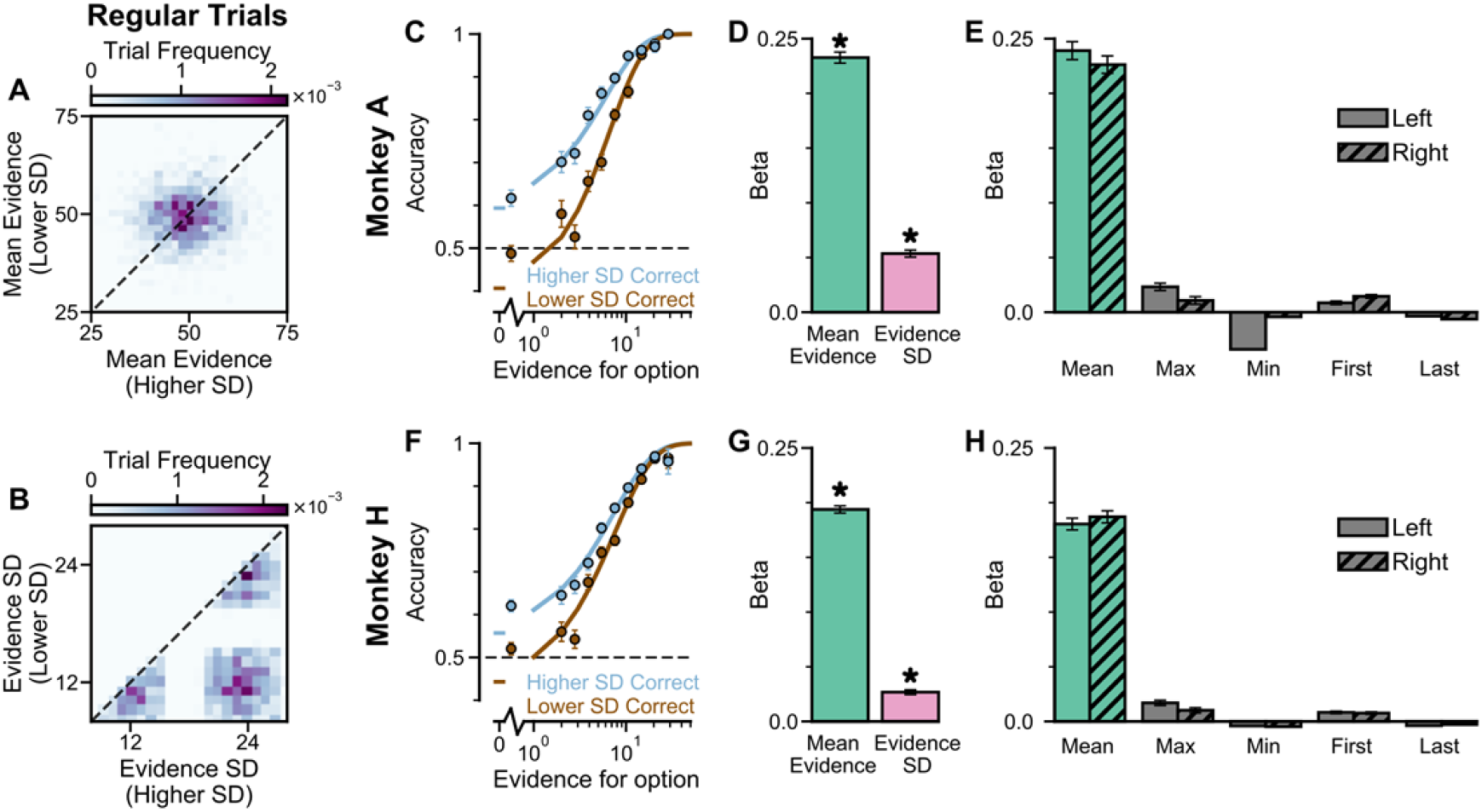
Extra information on Regular Trials: In the regular-trials, each of the two streams is randomly chosen to be either narrow (μ_N_ ∈ [47,53], σ_w_ = 12), or broad (μ_B_ ∈ [44,56], σ_B_ = 24), then divided into ‘Lower SD’ or ‘Higher SD’ options post-hoc, depending on the sampled standard deviation of evidence relative to the other option. (**A**) The distribution of the mean evidence of ‘Lower SD’ and ‘Higher SD’ streams, across all regular trials for both monkeys. (**B**) The distribution of the evidence variability of ‘Lower SD’ and ‘Higher SD’ streams, across all regular trials for both monkeys. (**C**) The psychometric function of Monkey A when either the ‘Lower SD’ (brown) or ‘Higher SD’ (blue) stream is correct. (**D**) A regression model using evidence mean and variability to predict the animals’ choices. Each regressor is the left-right difference of the mean and standard deviation of the evidence streams. This shows that both statistics are utilised by Monkey A to solve the task (Mean Evidence: t = 45.90, p < 10^−10^; Evidence Standard Deviation: t = 16.68, p < 10^−10^). (**E**) A regression model including the mean, maximum, minimum, first, and last evidence values of both the left and right streams as regressors, in order to evaluate the contribution of each quantity and the possibility that the monkey is utilising strategies alternative to evidence integration and pro-variance bias. Evidently, Monkey A mainly relies on temporal integration to solve the task, as indicated by a strong mean evidence coefficient in both regression models. See also **Supplementary Tables 1-3** for cross-validation analysis comparing regression models including various combinations of these predictors. (**F-H**) Same as (**C-E**) but for Monkey H. The statistics of the regression model in (**G**) are (Mean Evidence: t = 58.88, p < 10^−10^; Evidence Standard Deviation: t = 12.08, p < 10^−10^).

**Supplementary Figure 3:**
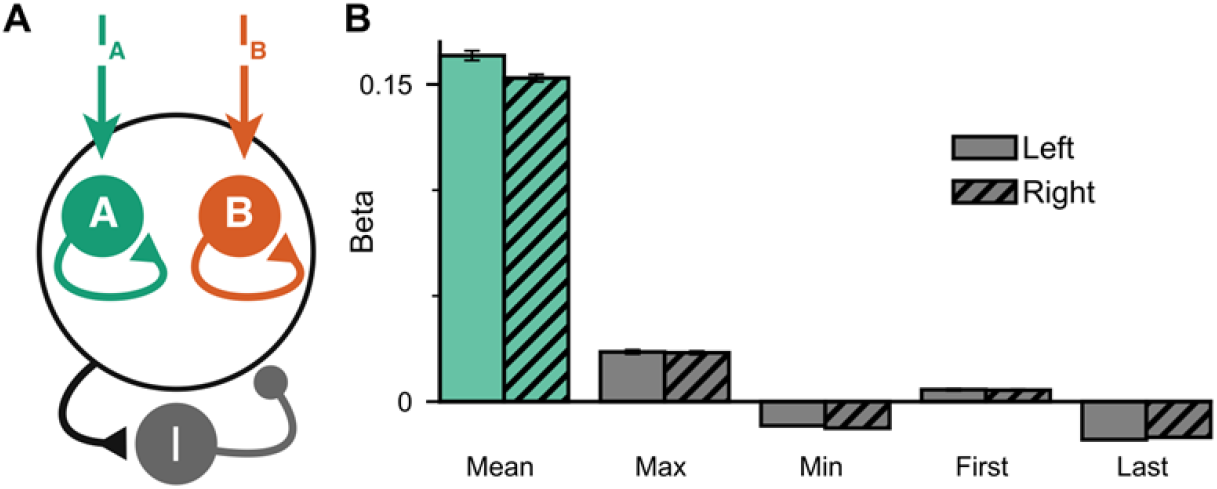
Extended regression results on the circuit model performance: (**A**) Circuit model schematic. The model consists of two excitatory populations which receive separate inputs, reflecting evidence for the two stimuli streams. Each population integrates evidence due to recurrent excitation, and competes with the other due to lateral inhibition. (**B**) Regression analysis of the regular trial circuit model data, using the mean, maximum, minimum, first, and last evidence values of both the left and right streams, in order to evaluate the possibility of decision-making strategies alternative to evidence integration and pro-variance bias. Similar to the monkeys, the circuit model mainly relies on mean evidence to solve the task. See also **Supplementary Tables 1-3** for cross-validation analysis comparing regression models including various combinations of these predictors.

**Supplementary Figure 4.**
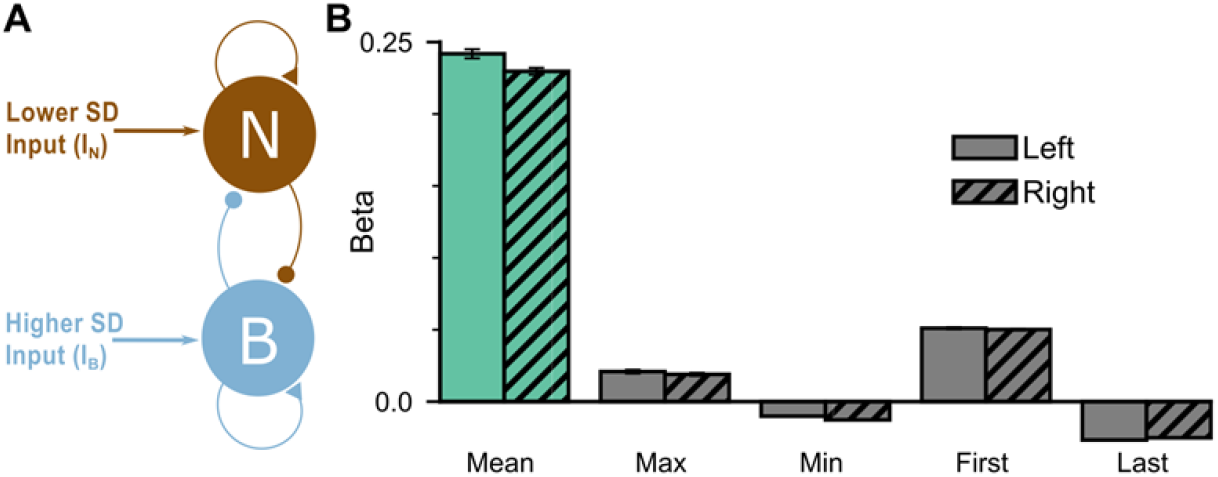
Extended regression results on the mean-field model performance: (**A**) The mean-field model consists of two variables which represent the accumulated evidence for the two choice options. The two variables demonstrate self-excitation and mutual inhibition. (**B**) Regression model on the regular trial model data, using the mean, maximum, minimum, first, and last evidence values of both the left and right streams, in order to evaluate the possibility of decision-making strategies alternative to evidence integration and pro-variance bias. Similar to the monkeys, the model mainly relies on mean evidence to solve the task.

**Supplementary Figure 5.**
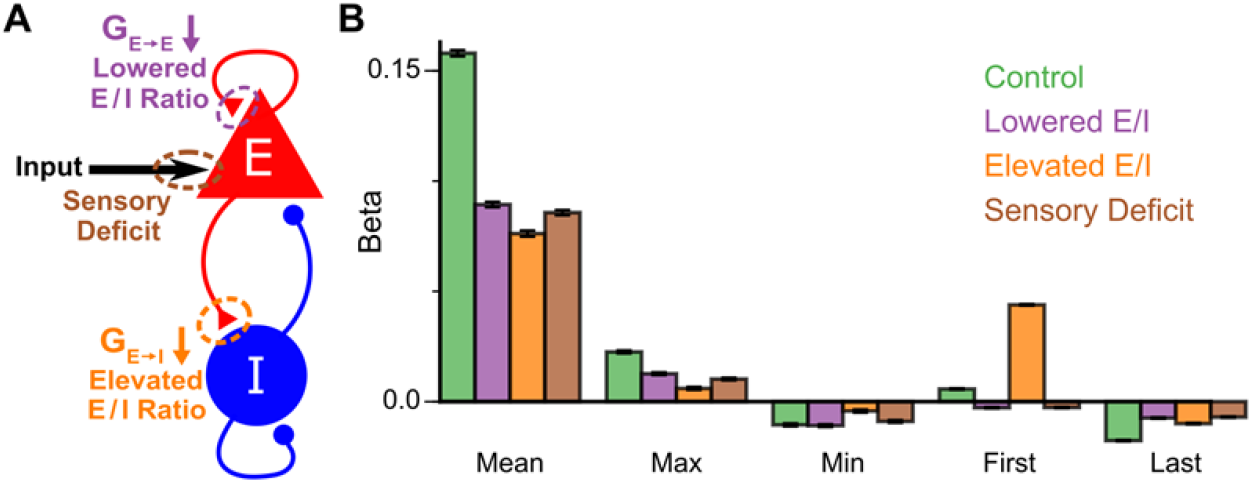
Extra information on Model excitation-inhibition (E/I) perturbations not influencing decisionmaking strategy: (**A**) Model perturbation schematic. Three potential perturbations are considered: lowered E/I (via NMDA-R hypofunction on excitatory pyramidal neurons), elevated E/I (via NMDA-R hypofunction on inhibitory interneurons), or sensory deficit (as weakened scaling of external inputs to stimuli evidence). (**B**) The regression model using mean, maximum, minimum, first, and last evidence values of each of the left and right streams as regressors, for the four models. Each bar shows the average of the left and right regressors of the corresponding variable. None of the perturbed models demonstrate a significant shift in decision-making strategies. The elevated E/I circuit has a larger first evidence regression coefficient, due to overemphasis of early stimuli (**Fig 7i**).

**Supplementary Figure 6.**
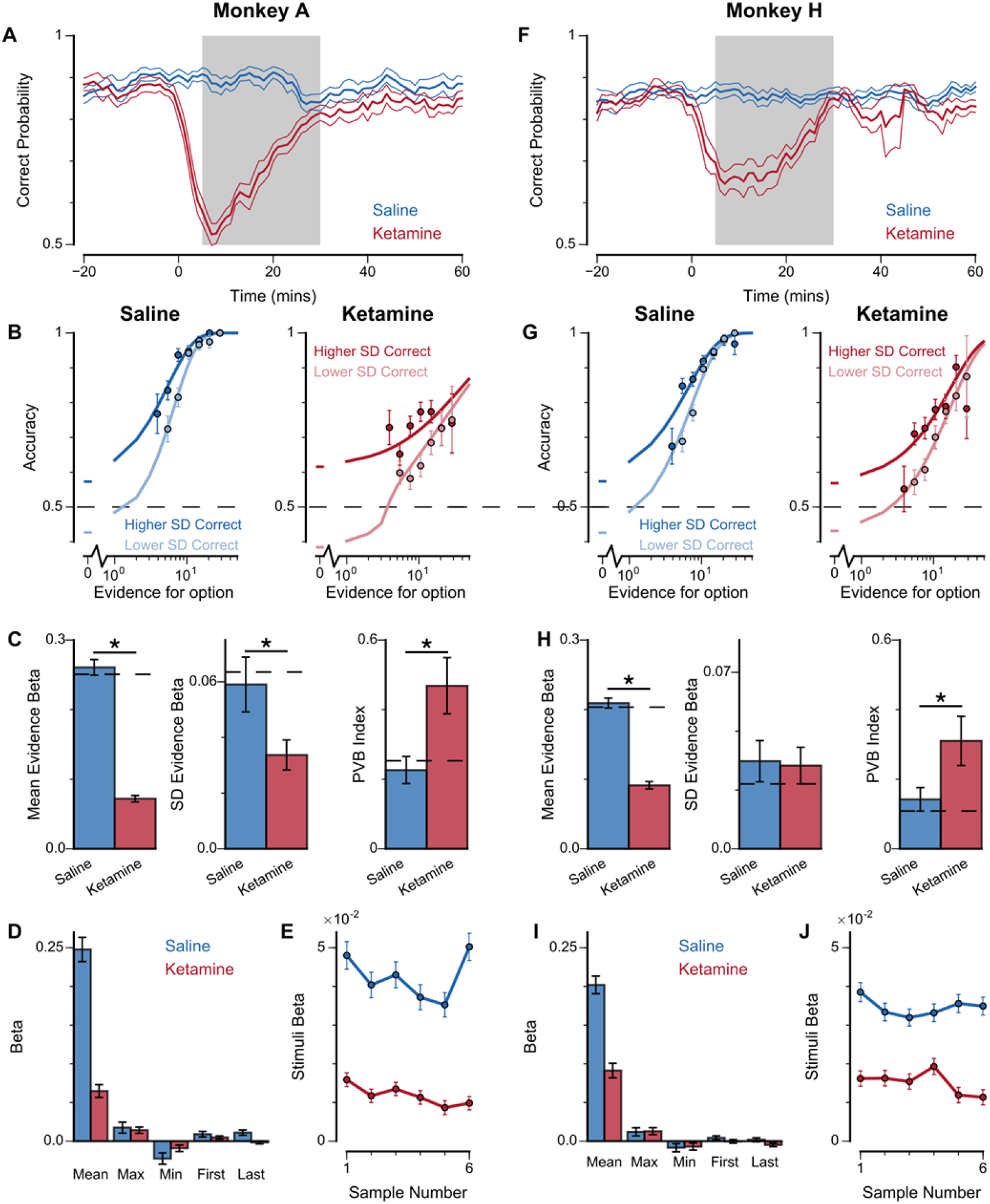
Extra information on ketamine experiments: (**A**) Mean percentage of correct choices across sessions made by Monkey A relative to the injection of ketamine (red) or saline (blue). (**B**) The psychometric function of Monkey A when either the ‘Lower SD’ or ‘Higher SD’ streams are correct with saline (**left**) or ketamine (**right**) injection. (**C**) Ketamine injection impairs the behaviour of Monkey A, in a manner consistent with the prediction of the lowered E/I circuit model. Dashed lines indicate pre-injection values in each plot. (**Left**) The regression coefficient for mean evidence, under injection of saline or ketamine. Ketamine significantly reduces the coefficient (permutation test p < 1×10^−6^), reflecting a drop in choice accuracy. (**Middle**) The regression coefficient for evidence standard deviation, under injection of saline or ketamine. Ketamine significantly reduces the coefficient (permutation test p = 4.98×10^−3^), but to a lesser extent than that of the mean evidence regression coefficient. (**Right**) Ketamine increases the PVB index (permutation test p = 1.16×10^−3^), consistent with the model prediction of the lowered E/I circuit. (**D**) The regression model using mean, maximum, minimum, first, and last evidence values of each of the left and right streams as regressors, under injection of saline or ketamine. Each bar shows the average of the left and right regressors of the corresponding variable. Ketamine injection does not alter decision-making strategies. (**E**) The regression weights of stimuli at different time-steps, for Monkey A with saline or ketamine injection. Ketamine injection lowers and flattens the curve of temporal weights, consistent with the lowered E/I circuit model. (**F-J**) Same as (**A-E**) but for Monkey H. (**E**) Ketamine significantly reduces the regression coefficient for mean evidence (permutation test p < 1×10^−6^), does not significantly reduce the regression coefficient for evidence standard deviation (permutation test p =0.871), and significantly increases the PVB index (permutation test p = 5.92×10^−3^).

## Supplementary Tables

**Supplementary Table 1:**
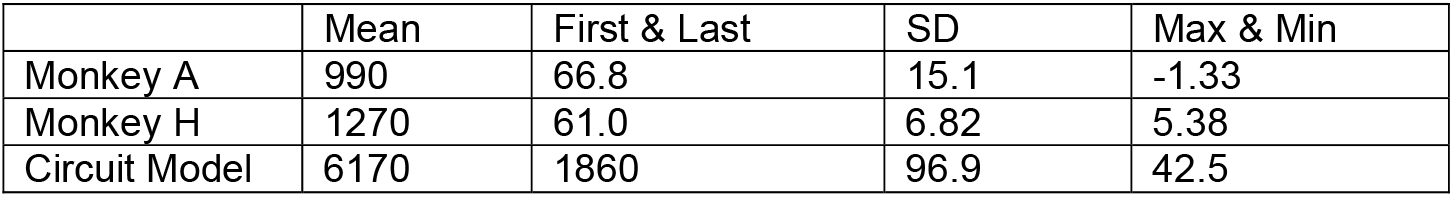
Difference in log-likelihood of Full regression model (mean, SD, max, min, first, last of evidence values; equation 6 in **Methods**) vs reduced model, for each monkey and the circuit model. Log-likelihood values were calculated using a cross-validation procedure (see **Methods**). Column label refers to the removed regressor. Positive values indicate the full regression model performs better. Values depend on the number of completed trials, which differed both between subjects and the circuit model. For both monkeys and the circuit model, mean evidence is clearly the most important driver of choice behaviour, followed by first and last evidence samples which reflects the primacy bias. Finally, evidence standard deviation (SD) has a stronger effect than maximum and minimum evidence samples (Max & Min).

|  | Mean | First & Last | SD | Max & Min |
| --- | --- | --- | --- | --- |
| Monkey A | 990 | 66.8 | 15.1 | -1.33 |
| Monkey H | 1270 | 61.0 | 6.82 | 5.38 |
| Circuit Model | 6170 | 1860 | 96.9 | 42.5 |

**Supplementary Table 2:** Difference in log-likelihood of regression models including either evidence standard deviation (SD) or both maximum and minimum evidence (Max & Min) as regressors, for each monkey and the circuit model. Log-likelihood values were calculated using a cross-validation procedure (see **Methods**). Column label refers to the regressors additional to either SD or Max & Min. Positive values indicate the regression model with SD performs better than that with Max & Min. Values depend on the number of completed trials, which differed both between subjects and the circuit model. Regardless of whether first and last evidence sample regressors are included, the models with standard deviation of evidence have higher log-likelihoods than the models with maximum and minimum evidence samples, indicating a better explanation of the data by standard deviation than by maximum and minimum evidence samples.

|  | Mean | Mean & First & Last |
| --- | --- | --- |
| Monkey A | 15.2 | 16.4 |
| Monkey H | 1.81 | 1.44 |
| Circuit Model | 54.5 | 56.7 |

**Supplementary Table 3:**
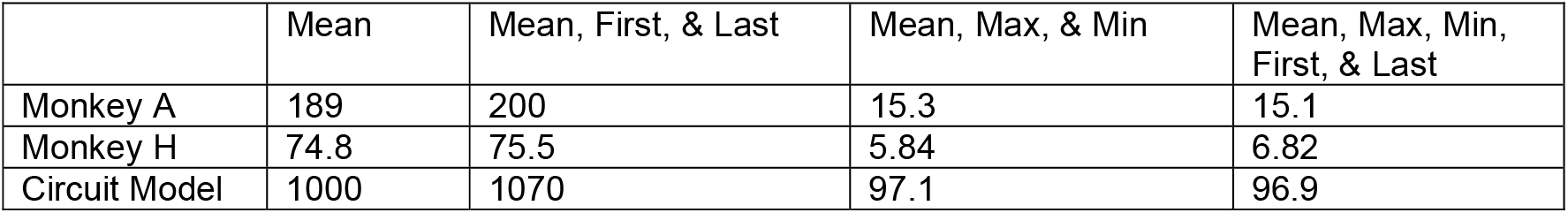
Increase in log-likelihood of various regression models (regressors in column labels) due to inclusion of evidence standard deviation as a regressor, for each monkey and the circuit model. Log-likelihood values were calculated using a cross-validation procedure (see **Methods**). Values depend on the number of completed trials, which differed both between subjects and the circuit model. Positive values across the table indicates the evidence standard deviation regressor robustly improves model performance for all models examined.

**Supplementary Table 4:** Difference in log-likelihood of regression models including either evidence standard deviation (SD) or both maximum and minimum evidence (Max & Min) as regressors, for each monkey with saline or ketamine injection. Loglikelihood values were calculated using a cross-validation procedure (see **Methods**). Column label refers to the regressors additional to either SD or Max & Min. Positive values indicate the regression model with SD performs better than that with Max & Min. Values depend on the number of completed trials, which differed across conditions. Regardless of whether first and last evidence sample regressors are included, the models with standard deviation of evidence have higher log-likelihoods than the models with maximum and minimum evidence samples, indicating a better explanation of the data by standard deviation than by maximum and minimum evidence samples. In particular, under ketamine injection, monkeys did not switch their strategy to primarily use maximum and minimum evidence samples (over standard deviation of evidence) to guide their choice.

|  | Mean | Mean & First & Last |
| --- | --- | --- |
| Monkey A Saline | 4.93 | 4.80 |
| Monkey A Ketamine | 2.91 | 2.69 |
| Monkey H Saline | 1.77 | 1.66 |
| Monkey H Ketamine | 2.50 | 2.23 |

